# Mechanistic multiscale modelling of energy metabolism in human astrocytes indicates morphological effects in Alzheimer’s Disease

**DOI:** 10.1101/2022.07.21.500921

**Authors:** Sofia Farina, Valérie Voorsluijs, Sonja Fixemer, David Bouvier, Susanne Claus, Stéphane P.A. Bordas, Alexander Skupin

**Affiliations:** Institute of Computational Engineering, University of Luxembourg, L-4364 Esch-sur-Alzette, Luxembourg; LCSB-Luxembourg Centre for Systems Biomedicine, University of Luxembourg, 7 Avenue des Hauts-Fourneaux, L-4362, Esch-sur-Alzette, Luxembourg; Department of Physics and Material Science, University of Luxembourg, L-1511 Luxembourg, Luxembourg; Luxembourg Center of Neuropathology (LCNP), Dudelange, Luxembourg; Laboratoire national de santé (LNS), National Center of Pathology (NCP), Dudelange, Luxem-bourg; Onera, 6 Chemin de la Vauve aux Granges, 91120, Palaiseau, France; Department of Neuroscience, University of California San Diego, USA

## Abstract

Astrocytes with their specialized morphology are essential for brain homeostasis as metabolic mediators between blood vessels and neurons. In neurodegenerative diseases such as Alzheimer’s disease (AD), astrocytes adopt reactive profiles with molecular and morphological changes that could lead to the impairment of their metabolic support and impact disease progres-sion. However, the underlying mechanisms how metabolic function of human astrocytes is impaired by their morphological changes in AD is still elusive. To address this challenge, we developed and applied a metabolic multiscale modelling approach integrating the dynamics of metabolic energy pathways and physiological astrocyte morphologies acquired in human AD and age-matched control brain samples. The results demonstrate that the complex cell shape and intracellular organization of energetic pathways determine the metabolic profile and support capacity of astrocytes in health and AD conditions. Thus, our mechanistic approach indicates the importance of spatial orchestration in metabolism and allows for the identification of protective mechanisms against disease-associated metabolic impairments.

---

The human brain is the organ with the highest energy demands required to sustain the high activity of neurons ^1^. Astrocytes are multitasking glial cells directly contributing to brain homeostasis and metabolism. By their complex architecture as star-like branched cells, they are intermediate structures sitting between neurons and their synapses, which they enwrap with their intricate processes, and the blood vessels, which they engulf with their endfeet. Based on this strategic positioning, astrocytes act as metabolic supporters providing energy in the form of lactate (LAC) to neurons and modulating their activity ^2, 3^. Astrocytes are also known to respond to brain “insults” and drastically change in many brain diseases such as Alzheimer’s disease (AD). In these situations, they engage reactive profiles with changes in morphology and in their molecular program ^4, 5^ like in AD where human astrocytes exhibit hypertrophy and overbranching ^6, 7^. In addition to morphological changes, AD-associated astrocytes also exhibit metabolic dysfunctions ^8–12^, altering their role as neuronal supporters, but the relation to morphology is not established.

The metabolic support function of astrocytes depends on sufficient LAC production and efficient LAC export at the perisynapses as energy substrate for neurons ^13^, and on sufficient availability of adenosine triphosphate (ATP) for its own metabolic sustainability ^14^ requiring an ATP : ADP ratio at least larger than one ^15^. Furthermore, physiological conditions for functional astrocytes are characterized by an approximate 10:1 ratio between LAC and pyruvate (PYR), the substrate for lactate production and mitochondrial activity, further indicating their metabolic support function ^16^. Hence, astrocytes have to keep a balance between a LAC based “altruistic” support mode and a more “egocentric” self-sustainability characterized by a high ATP : ADP ratio. The mechanistic relation between the observed disease-related modifications in morphology and metabolic dysfunctions are still to be characterized and whether morphology changes might represent a compensatory mechanism remains elusive.

Here, we develop a general interdisciplinary approach to systematically investigate the interplay between astrocytic morphology and energy metabolism in AD by a novel spatiotemporal *in silico* model that allows for physiologically realistic simulations by integrating complex morphologies obtained by high-resolution confocal microscopy and thereby addresses the impossibility of appropriate *in vivo* human astrocyte studies. Metabolic modelling has been extensively addressed in literature at different levels *via* detailed genome-scaled metabolic network models ^17^ or *via* targeted dynamic models ^18^, including astrocytic metabolism ^19–21^. All existing models neglect the spatial dimensions as they describe the metabolic processes through ordinary differential equations (ODEs). The underlying assumption that diffusion and reaction rates of metabolism are large enough to smear out spatial aspects are challenged by the complex morphology of astrocytes and an increasing amount of evidence for relocation of enzymes and other reaction site in different conditions ^22, 23^. To include spatial variations and geometric effects, we developed a metabolic model by means of a complex reaction-diffusion system (RDS) in realistic three-dimensional (3D) morphologies obtained from high-resolution confocal microscopy images of astrocytes in *post mortem* brain samples of AD patients and age-matched control subjects ^24^. The modelling framework incorporates the two essential astrocytic properties: 1) the main reactions of glucose metabolism are spatially localised to reflect the heterogeneous distribution of enzymes in the cell, and 2) the complex and context-dependent geometry of cells is directly incorporated from high-resolution microscopy. To address the resulting computational challenges in solving the corresponding partial equations of the RDS in realistic astrocytic morphologies with thin branches and regions of high curvature and kinks, we adapted our previous approach ^25^ utilizing the power of the cut finite element method (CUTFEM) ^26, 27^ to disentangle the complex astrocytic geometries from the mesh generation of finite-element methods and handle complex geometries as independently of the mesh as possible.

By this approach, our model paves the way to a more physiological modelling of the effect of astrocytic morphology in AD. Our framework is general and open-source and can be used for other cell types characterized by high-resolution imaging. For model establishment, we first performed simulations in simple two-dimensional (2D) geometries and studied how metabolic dynamics are affected by the spatial arrangement of reaction sites. The findings in 2D indicated the importance of the spatial component and the diffusion limitation that arise from the competition between the corresponding reaction centers for the metabolic substrates. Furthermore, the results highlighted the fundamental role of mitochondrial organization for the metabolic output of the system. Based on these insights, we subsequently investigated spatiotemporal metabolic dynamics in real 3D human astrocytic morphologies by our multiscale modelling approach and demonstrate the potential of our framework to study metabolic dysfunction in AD-related reactive morphology of astrocytes.

## Results

To investigate the potential mechanistic link between morphology and energy metabolic activity, our model describes glucose metabolism by five main metabolic pathways (Fig. 1**a**). We describe glycolysis *via* two subsequent pathways where the first represents the ATP consuming and the second one the ATP producing reactions. The first pathway is catalysed by a set of enzymes (hexokinase, phosphoglucose isomerase, phosphofructosekineaseand the fructose bisphosphate aldolase), which consume glucose (GLC) and ATP to produce ADP and glyceraldehyde 3-phosphate (GLY). In the following, we describe this pathway by a coarse-grained hexokinase (HXK) activity. In the second lumped reaction, these metabolites are transformed into ATP and pyruvate (PYR) by a second set of enzymes (glyeraldehyde phosphate dehydrogenase, phosphoglyerate kinease, phosophoglycerate mutase, enolase and the pyruvate kinase), which we describe by the overall activity of the pyruvate kinase (PYRK). The generated PYR is subsequently metabolised into LAC by the lactate dehydrogenase (LDH) or used by mitochondrial metabolism to generate ATP. The mitochondrial metabolic activity of the Krebs cycle and oxidative phosphorylation is described by the coarse-grained effective reaction Mito. Finally, another effective reaction (act) accounts for various ATP-consuming processes associated to cellular activity. These metabolic pathways are put into a spatial context by distributing the corresponding reaction centers into a spatial domain.

**Fig.1.**
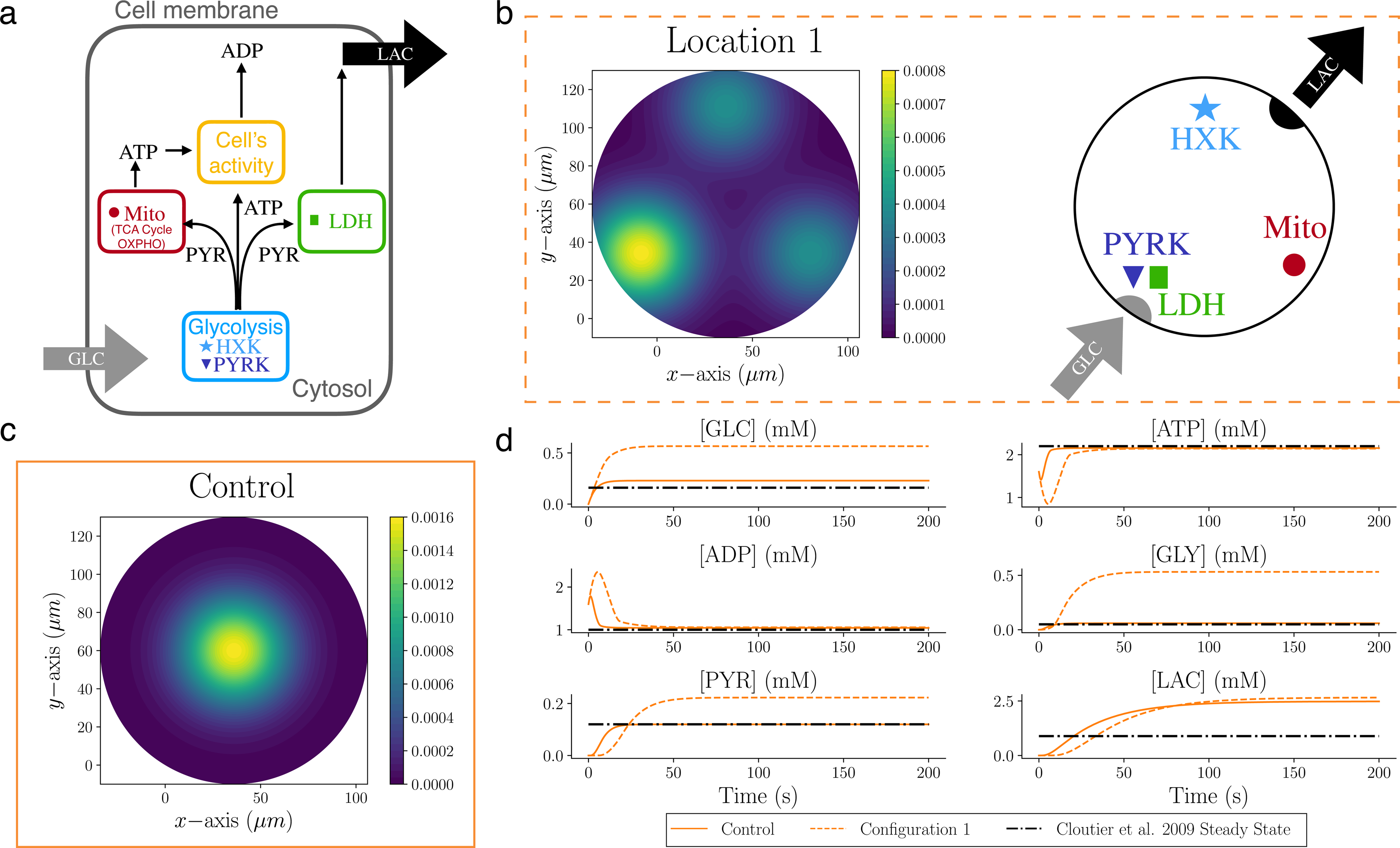
Spatial arrangement of metabolism has an impact on cellular metabolite concentrations. **a** GLC enters the cytosol of the cell and takes part in glycolysis whose effective kinetics is captured by the two reactions HXK and PYRK. The products of glycolysis are subsequently consumed by the LDH reaction for generating LAC, by the act reaction describing ATP consumption due to cellular activity, and by mitochondria where the effective reaction Mito produces ATP from PYR through the Krebs cycle and oxidative phosphorylation. **b** (left) Generic configuration to investigate the effect of metabolite transport on the output of metabolism in a 2D circular domain. The color map highlights the position of the reaction sites, which are located on the vertices of an equilateral triangle. Two reaction sites are colocalised at the bottom left corner. (Right) Position of the reaction sites in Location 1: HXK on the top close to the efflux of LAC, PYRK and LDH colocalised close to the GLC influx, and Mito on the last vertex. **c** Control scenario: all reaction sites are located in the center. **d** Dynamics of the average concentration of each species in Location 1 and control cases, compared with the steady state values from Cloutier *et al.*.

### Metabolic dynamics and reaction sites competition in 2D domains

For model establishment and calibration, we first analyzed the effect of different spatial arrangements of reaction sites on the metabolic profile in simple 2D geometries. For this, we considered a circular domain and compared different configurations of reaction locations. The diameter of the circular domain was set to 140 *µ*m as an average diameter that contains a full astrocyte ^28^. To reflect the metabolic flux from the endfeet towards the perisynapses at the neurons’ locations, we placed the entry of GLC and the exit of LAC at opposing sides of the circle (Fig. 1**b**) where the subregions are defined as the intersection of a circle with a radius of 10 *µ*m and centers are located at the origin for GLC and the antipodal point for LAC .

In this simplified setup, we first assumed that a given reaction occurs around a single location with a spatial extent of a Gaussian distribution with a width of *σ* = 20.0 *µ*m. As a control case, all four reactions were located in the center of the circle as shown in Fig. 1**c**, mimicking a well-stirred condition. In a more complex enzyme arrangement, we located the four reactions on the vertices of an equilateral triangle inscribed inside the circle: one reaction is placed on the top vertex close to the LAC exit, one reaction on the bottom right vertex and two reactions are placed on the bottom left vertex. An example is shown as “Location 1” in Fig. 1**b**, where PYRK and LDH are placed on the top of each other, while HXK and Mito are on the top and bottom right vertex, respectively.

The resulting dynamics of the Location 1 and the control setup are shown in Fig. 1**d** where the average concentration dynamics inside the domain for each involved metabolite is shown. As a reference, we also plotted the steady state concentrations from Cloutier *et al.* ^20^, which was used to calibrate parameters of our model for which no literature information was available (see Table 1). As expected, the control configuration leads to steady-state concentrations in agreement with Cloutier *et al.* during an equilibrium period of *≈* 50s, with an exception for LAC which exhibits an almost doubled level. By contrast, the arrangement of the enzyme sites in spatially distributed configurations such as Location 1 affects the metabolite levels of interest. For example, the steady state corresponding to Location 1 is characterized by concentrations of GLC, GLY, PYR and LAC that are approximately four, ten, two and three times higher compared to the well-mixed condition described by ODEs in Cloutier *et al.*, respectively.

**Table 1:**
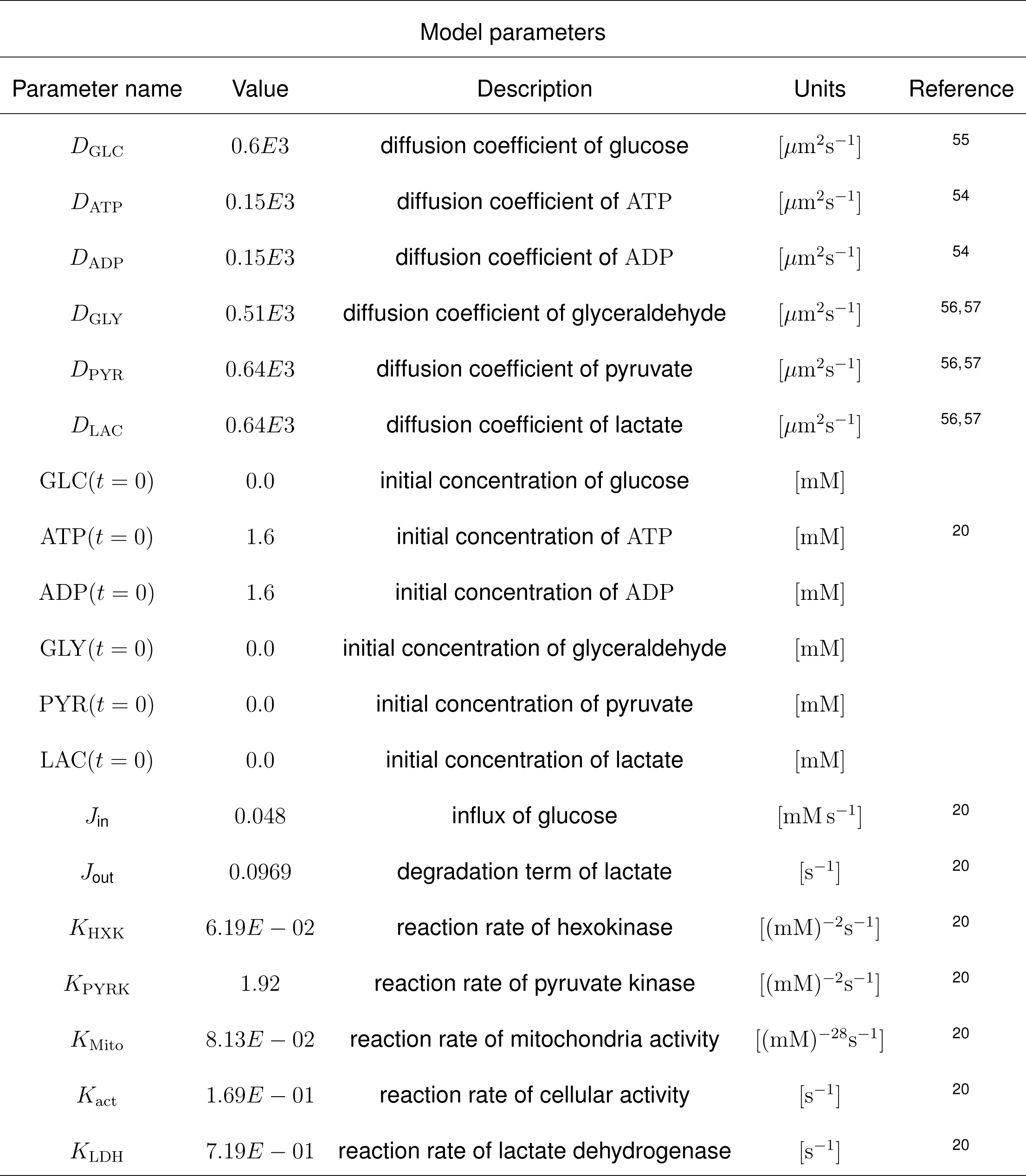
Model parameters

The steady state solutions of Location 1 indicates the necessity of the species to diffuse inside the domain and reach the corresponding enzyme sites: GLC needs to diffuse into the other part of the domain to act as substrate for HXK, and the produced GLY needs to reach the PYRK to be metabolized into PYR. The reactions are thereby diffusion-limited and the system reaches the steady state before consuming more GLY. Finally, the increased LAC level for Location 1 in relation to the control case is caused by the co-localization of PYRK and LDH where produced PYR is directly metabolized into LAC whereas in the control case Mito and PYRK compete for PYR as substrate.

To investigate systematically the effect of co-localisation and/or proximity of reaction centers to GLC influx or LAC efflux, we considered all possible location configurations for the four reactions on the vertices of the triangle (Fig. 1**b**). Considering the colocalisation of two reactions in the left-bottom vertex, leads to twelve possible location configurations (Figs. 2**a** and 2**c**). As a first attempt to address slightly more complex morphologies, we studied the twelve locations within a two-dimensional star shape (Fig. 2**b**) as a simplified version of an astrocyte. This setup allows for comparable results between the two domains, since molecules have to pass similar distances between the subregions where GLC enters and the subregion where LAC is exported. Reaction sites were located analogously at the three vertices of an equilateral triangle within the star. As in the circular setup, we placed two reaction sites colocalised closer to the influx of GLC, and one reaction site at each of the remaining vertices.

**Fig.2.**
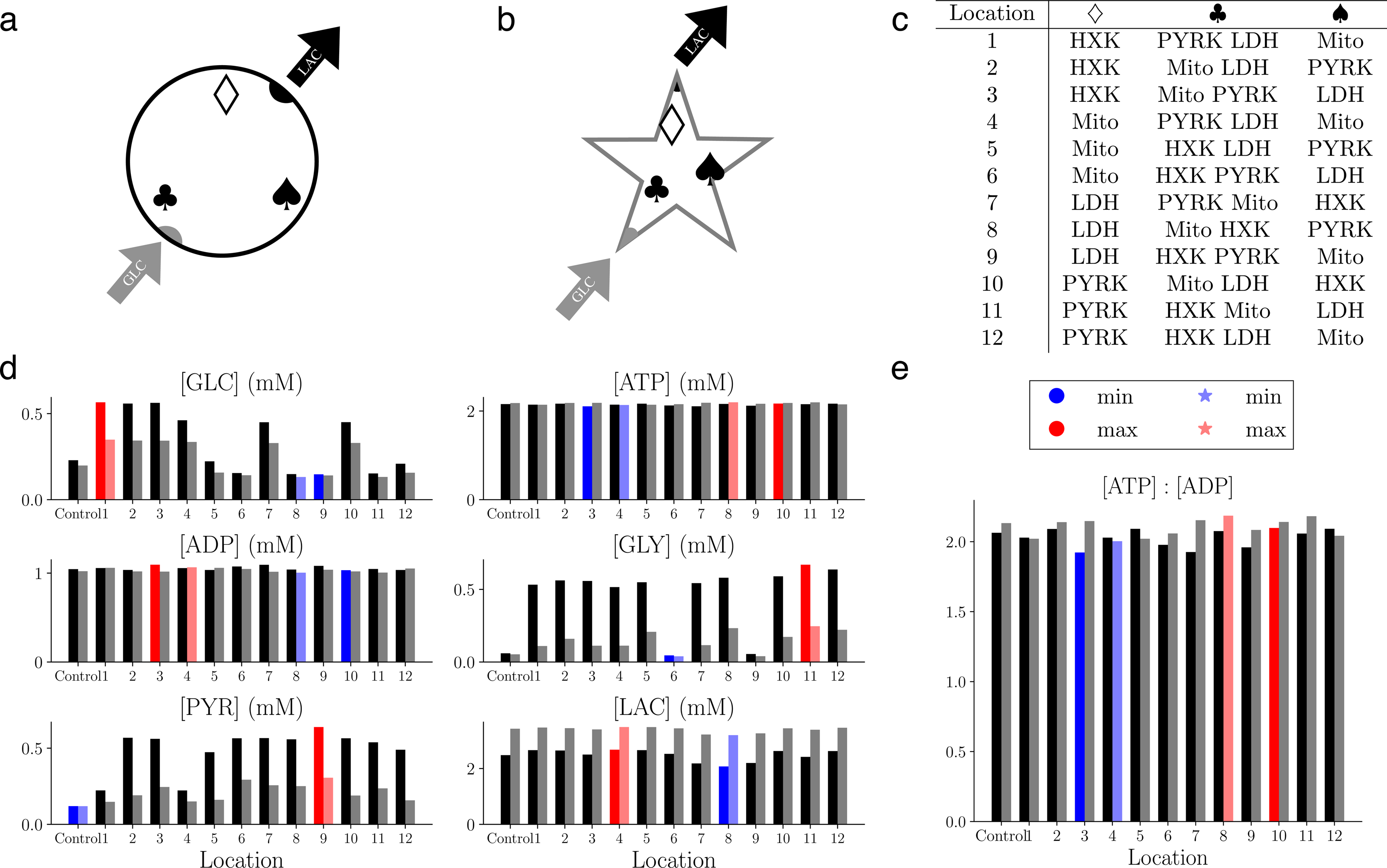
Spatial organization and competition between reaction sites affect the metabolic activity of the system. **a** Spatial setting of simulations performed in a 2D circle. GLC enters along one side, and on the diametrically opposite side, LAC is exported. Each of the three symbols is associated to one (diamond and spade) or two (club) reactions. **b** Spatial setting of simulations performed in the star shape. The reaction sites are located analogously to the circle domain with the same distance between the GLC entry vertex and the LAC efflux/degradation. **c** Table of the 12 possible configurations corresponding to the allocation of one reaction site to diamond and spade vertices, and two colocalised reaction sites at the club vertex. **d** Spatially averaged steady-state concentrations of each species for the circle (left) and the star (right) **e** Spatially averaged steady state ATP : ADP ratio for simulations in a circular (left) and star-like geometry (right).

Fig. 2**d** shows the steady-state and spatially averaged concentration of each species of interest for the twelve possible configurations of the circular (left columns) and the star domain (right columns) where the maximum and minimum values for each species are highlighted in red and blue, respectively. Simulations performed in both domains exhibit similar trends. The species that are affected the most by the different spatial arrangement are GLC, GLY and PYR for the co-localisation of the reaction sites HXK-PYRK (Location 6 and 9) and PYRK-LDH (Location 1 and 4) which led to low level of GLY or PYR, respectively. In the control case, where all the reaction sites overlap in the center of the domain, the system is more efficient with low levels of GLC, GLY and PYR, and a medium value of LAC. Although LAC shows differences depending on the location of the reaction sites, the changes are less significant due to the efflux which reduces LAC steady state concentrations. Interestingly, the star-shape domain exhibits the highest values of LAC pointing to the importance of morphologies with branches and higher complexity. Since cellular activity is assumed to occur homogeneously inside the domains, variability in ATP and ADP levels across the setups are rather small confirmed by the [ATP] : [ADP] ratio with a variance between all the simulations of 0.005 (mM^2^) (Fig. 2**e**).

Overall, these *in silico* experiments emphasize the variable output of the metabolic RDS as a function of the intracellular spatial organization of reaction sites. To further investigate this effect, we next modelled the effect of enzyme distributions in more detail.

### Uniform and polarised distribution of reaction sites in a rectangular domain

Based on the establishment of the spatiotemporal metabolic model for one reaction center for each pathway reaction, we next explored the effect of inhomogeneous distributions of reaction centers on the metabolic state of the cell. For this purpose, we considered for each metabolic reaction ten distinct reaction sites with a smaller spatial extent (*σ* = 1.0 *µ*m), while conserving the overall metabolically activity. To mimic the morphology of astrocytic branches, the shape of the RDS domain was chosen as a two-dimensional rectangle of dimension [0*, l*] *×* [0*, L*], with width *l* = 4 *µ*m and a length *L* = 140 *µ*m where GLC enters from the bottom left corner of the rectangle (origin) and LAC exits from the top right corner. We considered two types of cellular organisation: one where the reaction sites are uniformly distributed inside the domain and the extreme opposite setting of a polarised cell where some reactions occur predominantly at one of the extremities of the cell. To ensure robustness of the findings, the two settings were compared by ensemble simulations of 200 distinct realizations of each setting. For the uniform cells, the coordinates of the 10 reaction sites of each type were randomly selected from a uniform distribution that covers the rectangular domain. Realizations of polarised cells were generated either by normal distributions (*N* (*m, σ^1^*), where *m* and *σ^1^* denote the mean and standard deviation, respectively) or by log-normal (log *N* (*m, σ^1^*)) distributions. Fig. 3**a** shows the position of the reaction sites along the *y* coordinate of the 200 realisations and Fig. 3**b** exemplifies enzyme distributions for a given cell for each setting. The different strategies for polarised cells lead to a certain probability for mitochondria localization in the upper part of the domain for the “Polarised” configuration but not for the “Polarised log *N* (2)” configuration (Fig. 3**a**). These settings allow for investigating the competition between the Mito and LDH reactions for their shared substrate PYR (Supplementary Note 1).

**Fig.3.**
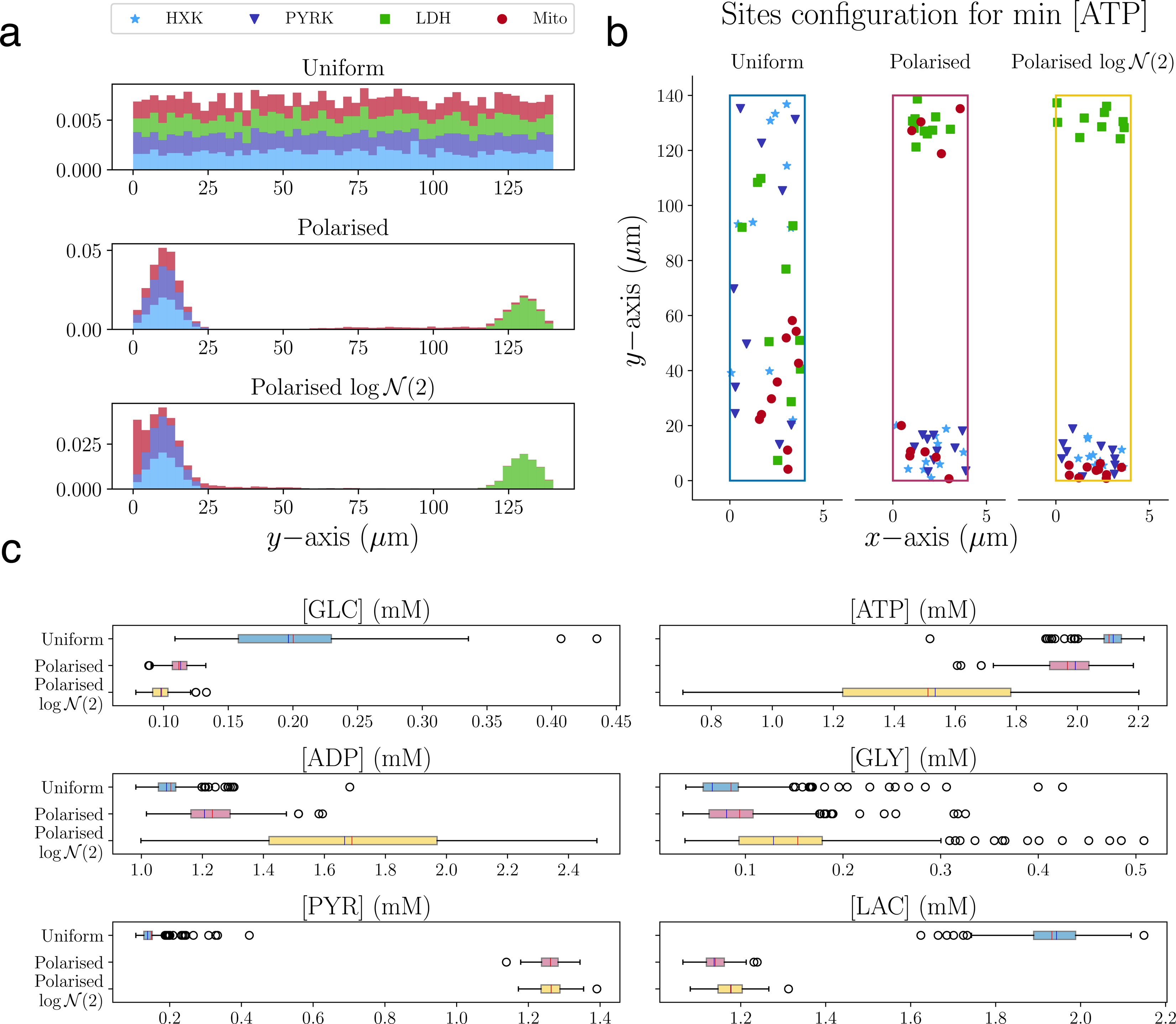
The steady-state level of metabolites is affected by the polarised distribution of enzymes within cells. **a** Distribution of the enzymes along the *y*-axis of the rectangular domain: HXK in light blue, PYRK in dark blue, LDH in green and Mito in dark red (top panel). In the Uniform setting, the reacting sites are uniformly distributed along the *y*-axis. In the Polarised settings, HXK, PYRK and LDH are spread unevenly over the domain with the first two located close to the origin and the latest close to the top of the domain. Mito reaction sites are distributed in the following way: 6 of them are normally distributed and colocated in the same area as HXK and PYRK, and 4 of them are uniformly located in the upper part of the domain (middle panel). In the Polarised log *N* (2) setting, mitochondria are located in the domain according to a log-normal distribution (bottom panel). **b** Examples of Uniform, Polarised and Polarised log *N* (2) distributions for the less energized cell where mitochondrial production is the most affected by polarisation. **c** Box plot of the average steady-state concentration of each species for the Uniform, Polarised and Polarised log *N* (2) distributions. (The mean and median of each box is signed in red and blue, respectively.)

To assess the effect of the different spatial arrangements, the steady state concentration of the 200 realizations, for the three different configurations, were compared statistically (Fig. 3**c**) including T-test and Wilcoxon-Mann-Whitney with Holm-Bonferroni compensation (Table 2 and Supplementary Note 2). In general, the polarised cells consume more GLC than the uniform distributed ones, which is consistent with the fact that the reaction HXK is closer to the influx. GLY is present at a very low level for all configurations as also shown in the significance test. Interestingly, PYR and LAC differ strongly in polarised cells compared to the uniform setting with a higher level in PYR caused by faster metabolizing of GLC by the HXK and subsequent PYRK reactions. On the other hand, LAC levels are higher for the uniform cells since in polarised cells PYR reaches the more distant LDH reaction only by the amount which has not been consumed by the closer located Mito reactions. The resulting LAC : PYR concentration ratio for the cells with uniformly distributed enzymes cells respect the physiological constraints, whereas polarised cells exhibit ratios below one indicating an unphysiological or diseased state.

**Table 2:**
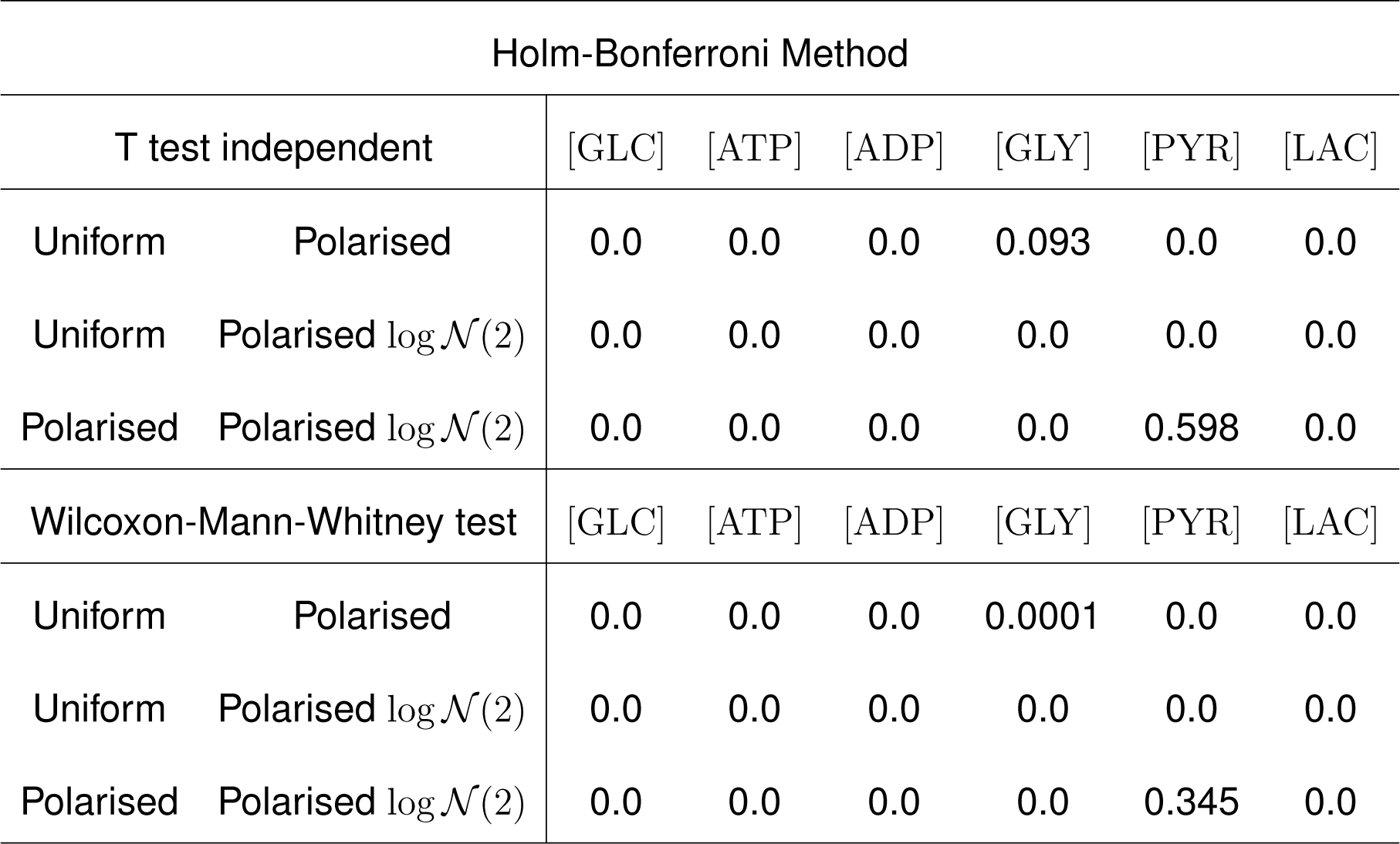
*p*−values of the significance tests for Experiment 2. We used multiple comparison Holm-Bonferroni Method on a parametric independent T-test and a non-parametric Wilcoxon-Mann-Whitney test with significance threshold of p−value < 0.05.

| Holm-Bonferroni Method |  |  |  |  |  |  |  |
| --- | --- | --- | --- | --- | --- | --- | --- |
| T test independent |  | [GLC] | [ATP] | [ADP] | [GLY] | [PYR] | [LAC] |
| Uniform | Polarised | 0.0 | 0.0 | 0.0 | 0.093 | 0.0 | 0.0 |
| Uniform | Polarised $\log \mathcal{N}(2)$ | 0.0 | 0.0 | 0.0 | 0.0 | 0.0 | 0.0 |
| Polarised | Polarised $\log \mathcal{N}(2)$ | 0.0 | 0.0 | 0.0 | 0.0 | 0.598 | 0.0 |
| Wilcoxon-Mann-Whitney test |  | [GLC] | [ATP] | [ADP] | [GLY] | [PYR] | [LAC] |
| Uniform | Polarised | 0.0 | 0.0 | 0.0 | 0.0001 | 0.0 | 0.0 |
| Uniform | Polarised $\log \mathcal{N}(2)$ | 0.0 | 0.0 | 0.0 | 0.0 | 0.0 | 0.0 |
| Polarised | Polarised $\log \mathcal{N}(2)$ | 0.0 | 0.0 | 0.0 | 0.0 | 0.345 | 0.0 |

The corresponding ATP and ADP concentrations show a rather low variability for the uniform configuration with higher ATP and lower ADP concentrations (Fig. 3**c**) compared to the polarised cells. Interestingly, the Polarised log *N* (2) configuration exhibits a very wide range for both concentrations with significantly different average values also in comparison with the Polarised configuration indicating the importance of mitochondrial distribution. Thus, the ATP : ADP ratio for the three cellular configurations (Fig. 4**a**) confirms that the Polarised log *N* (2) realisations cover an ATP : ADP ratio range from unhealthy (ratio *<* 1) to healthy (ratio *≥* 1).

**Fig.4.**
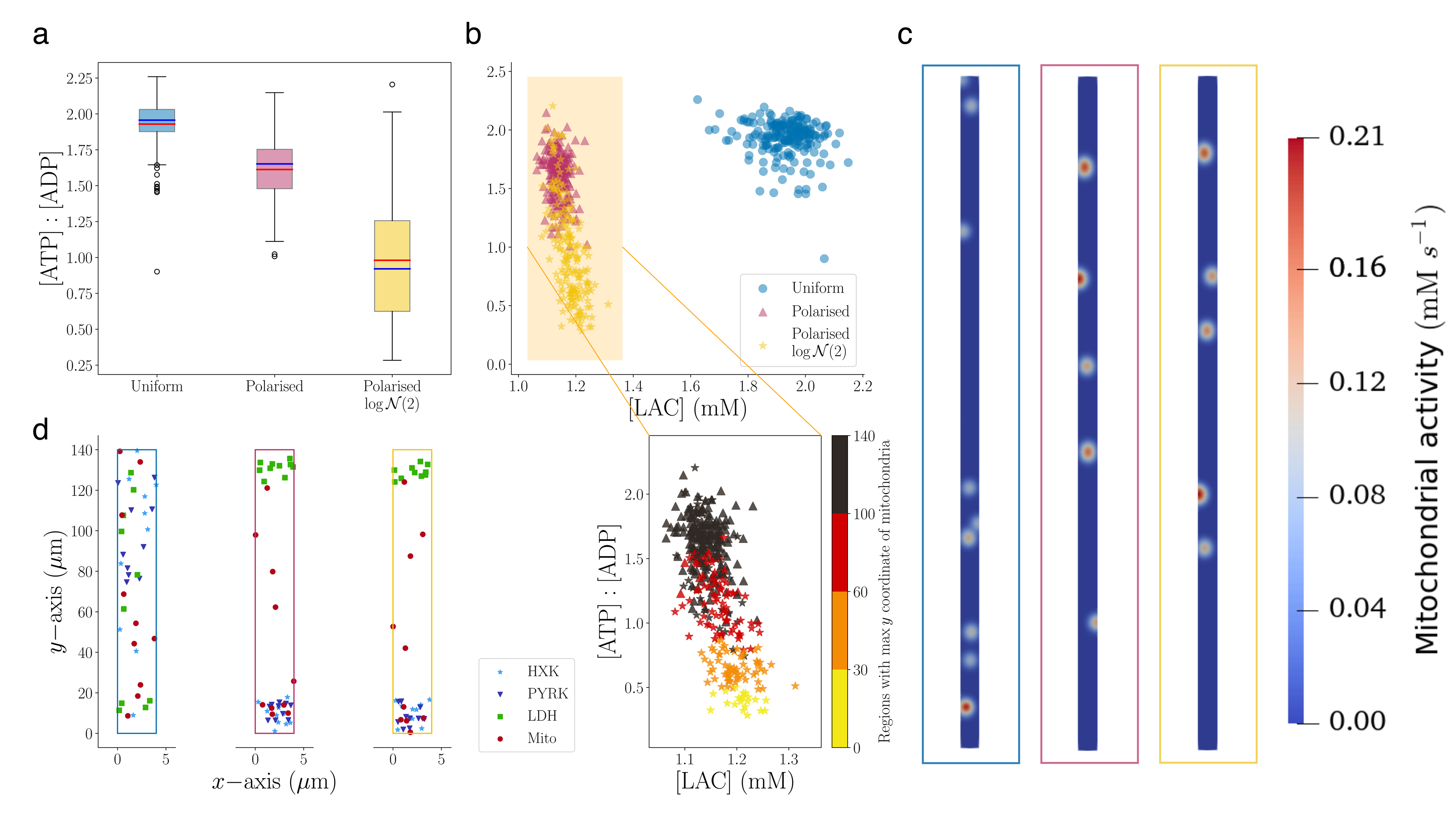
Mitochondrial distribution determine [ATP] : [ADP] ratio and thereby energetic states of cells. **a** Box plot of the final average values of the [ATP] : [ADP] for Uniform, Polarised and Polarised log *N* (2) (Mean in red and median in blue). **b** (top) scatter plot of ratio against LAC final average values. There are two distinct clusters between the Polarised cells and the uniformly distributed ones. (bottom) Zoom on the ratio against LAC for Polarised cells colored based on the region where we can find the mitochondria with the highest *y*-coordinate. Interestingly if the enzymes are well distributed inside the domain, so if there is at least one mitochondria with a *y*-coordinate larger than 100 (black), the ratio value is higher. On the other hand, if the mitochondria are all located within the first or second region (yellow and orange), with *y*-coordinate lower than 60, the cell is in unhealthy status. **c** Mitochondria activation of the configurations with maximum ATP production for the three type of cells at steady state **d** Reaction sites setting for the maximum level of ATP for the three distributions.

The impact of the configurations on the metabolic activity and in particular with a focus on the “altruistic” behaviour producing more LAC or an “egocentric” strategy producing more ATP, can be visualized by the relationship between LAC and the ATP : ADP ratio (Fig. 4**b**). We found two distinct clusters formed by uniform and polarised cells, where the uniform cells display a co-existing egoistic and altruistic mode characterized by high ATP and LAC concentrations for self-sustainability and neuronal support. Indeed, the correlation between the variables, the ratio and LAC, is only slightly negative for the uniform cells (*−*0.19), whereas the group of polarised cells exhibits stronger negative correlation (*−*0.65) indicating that high values of one quantity lead to low values of the other. “Polarised” cells are located on the top of the cluster and the “Polarised log *N* (2)” cells are predominantly in the lower part of the cluster characterized by a lower ATP : ADP-ratio and slightly higher LAC concentrations (Fig. 4**b**). This difference indicates the importance of mitochondria localization as shown by the color-indicated classification of the vertical arrangement of mitochondria. We colored the metabolic profile of each realisation based on the highest *y−*coordinate of Mito sites (*y*_max_): yellow, orange, red or black, if *y*_max_ *<* 30, 30 *< y*_max_ *<* 60, 60 *< y*_max_ *<* 100 or *y*_max_ *>* 100, respectively. This analysis highlights that the lowest level of ATP coincides with realisations where all mitochondria are grouped in the lower region. By contrast, simulations with a high energetic profile correspond to arrangements where mitochondria are distributed throughout the whole rectangular shape. The spatial arrangements and the corresponding mitochondrial activation that describe the most energized cell for each configuration are shown in Figs. 4**c**-**d** and confirm the necessity of mitochondria to be well-distributed in the whole domain to sustain high ATP levels. Comparing the mitochondrial activity at steady state for each of these cells, we notice low activity in the lower part of the polarised cells indicating the co-localization of Mito and PYRK leads to substrate competition. This suggests that PYRK inhibits the mitochondrial activity. On the other hand, the cellular arrangements presented in Fig. 3**b** produce the less energised cells with minimum ATP and all mitochondria gathered in clusters.

Overall, this analysis demonstrates the impact of the interplay between spatial enzyme orchestration and morphology on the metabolic profile of cells. Our finding highlights that different cellular organization leads to different steady state concentrations which might be linked to potential disease of cells.

### Morphological effects on metabolic activity of human astrocytes in health and AD

Finally, we extend our work to 3D reconstructions of human astrocytes acquired from GFAP-immunostained *post-mortem* brain samples from age-matched control subjects (Figs.5**a**-**c**) and AD patients (Figs.5**d-f**). The 3D confocal images of the astrocytes were acquired in the CA1 subregion of the hippocampus (Figs. 5**a** and 5**d**). Given the typical *post-mortem* nature of such brain samples, the dynamical consequences of the morphology for metabolic profiles can be only assessed by an appropriate *in silico* strategy. The respective segmentations of the prototypical astrocytes (Figs. 5**b** and 5**e**) reveal significant differences in the volume and morphological diversity of the two cells: the reactive AD astrocyte exhibits hypertrophy, proliferation of branches and coverage of wider spatial domains in comparison with the less complex shape of the control astrocyte (Figs. 5**g** and 5**h**). Based on mitochondria staining and segmentation (Figs. 5a-b and 5**d-e**), a realistic spatial arrangement of mitochondria is implemented in the multiscale model (Figs. 5**c** and 5**f**). The presence of regions with different mitochondrial density is respected by tuning the center positions and variances of the Mito spatial reaction rates (Figs. 5**c** and 5**f**). The minimum variance is set to 1.0 *µ*m and we scale accordingly the size of the other regions with a maximum of 2.0 *µ*m. The number of reaction sites for the other reactions is set according to the amount of mitochondria selected from postprocessing, 97 for the control and 140 for the reactive astrocyte (Fig. 5**i**). For each Mito reaction site, we located a HXK site close by in agreement with the observed relationships between these two enzymes ^29, 30^. The reaction sites of PYRK and LDH are taken from a uniform distribution defined in the three dimensional box containing the astrocyte. The locations of the reaction sites for the simulations inside the control and AD reactive astrocyte are shown in Figs. 6**a** and 6**c** together with the assumed endfeet for GLC influx and the subregions at the perisynapses for LAC export into the extracellular space (Figs. 6**a-c**).

**Fig.5.**
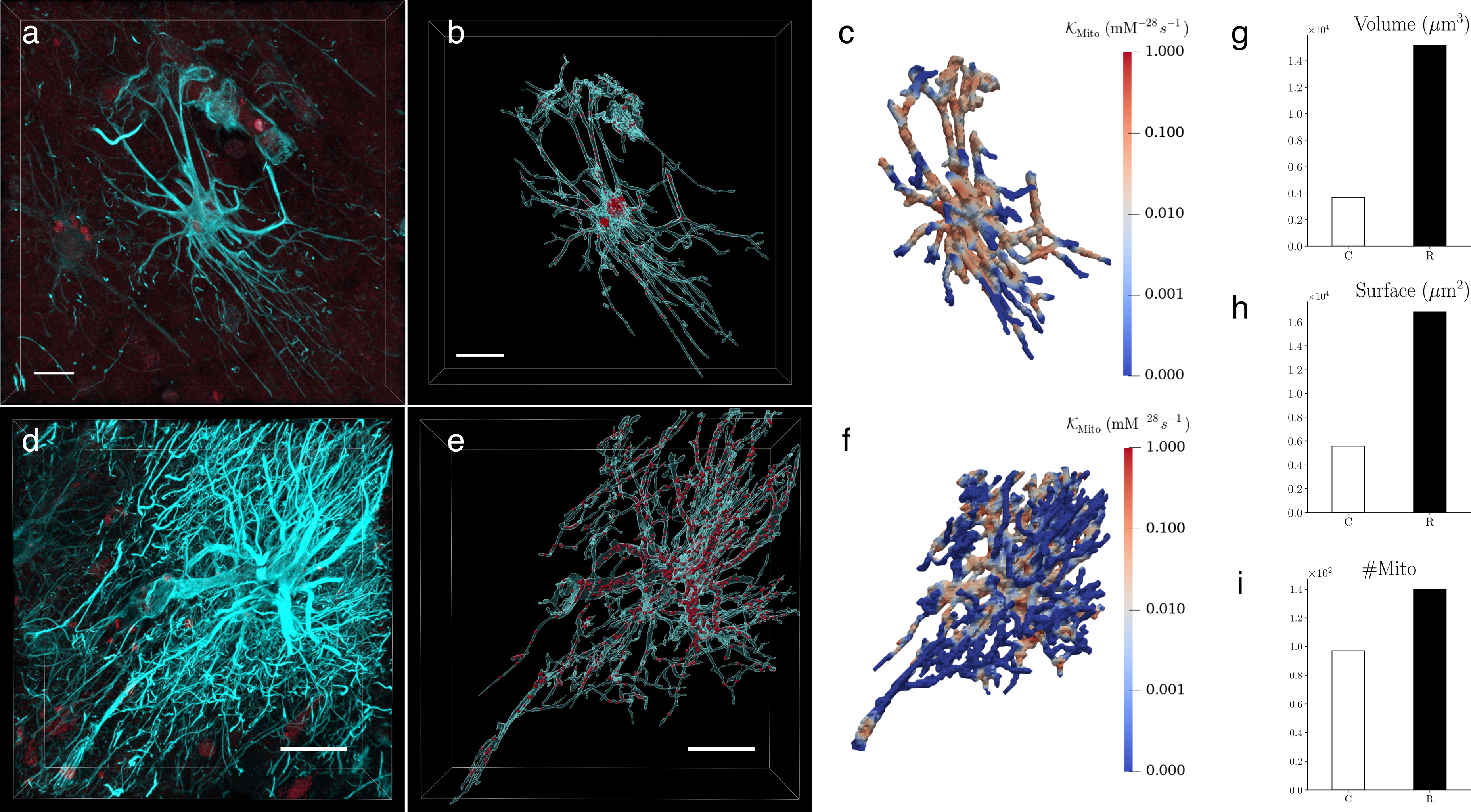
Human hippocampal astrocytes from an age-matched control subject and AD patient: from microscopy image to 3D simulation setting. High-resolution confocal microscopy images **a** from an age-matched control subject and **d** from an AD patient were obtained from 50 *−* 100 *µ*m brain sections that were immunostained against GFAP (cyan) to visualize astrocyte cytoskeletal morphology, and against TUFM (dark red) to reveal mitochondria in the hippocampus. Using Imaris 9.6.0 **b** and **e**, astrocyte 3D morphology was segmented using the surface tool and mitochondria were labelled with the spots tool for the astrocytes in both conditions. Finally, based on the segmentation, we created the domains for our simulations and we selected the locations with higher density to define the Mito reactions. **c** and **f** show the spatial reaction rates *K*_Mito_ describing the mitochondria activity inside the cells. In the bar charts **g-i**, we compared the cell volumes – 3673 *µ*m^3^ for the control astrocyte (C) and 15161 *µ*m^3^ for the reactive (R), cell surfaces – 5569 *µ*m^2^ for C and 16854 *µ*m^2^ for R, and the number of mitochondria activity centers – 97 for C and 140 for R computed in **c** and **f**. Scalebars: **a**-**b** 15 *µ*m, **d**-**e** 30 *µ*m.

As a first analysis, we ran three baseline simulations based on the physiological parameters (Table 1) with one simulation inside the protoplasmic control morphology (C) (Fig. 6**a**), one within the same morphology but with a polarised distribution of reaction centres (P) (Fig. 6**b**) and one inside the reactive astrocyte (R) (Fig. 6**c**, more details are given in Supplementary Note 3). The resulting dynamics of these baseline simulations (Fig. 6**d**) are in good agreement with the investigation of enzyme distributions in the 2D domains where scenarios C and R resembles properties of the uniform distributed cell and P corresponds to the polarised cell (Fig. 3**b**). However, the average LAC concentration is higher than expected for P and lower than expected for C and R. Also the PYR concentration is closer to that of the uniform setting for the P configuration. While the concentration values of C and R are on average very close (Fig. 6**b**), smaller differences are visible mostly in GLC and LAC, and attributed to the effect of the morphological differences and reaction site configurations.

**Fig.6.**
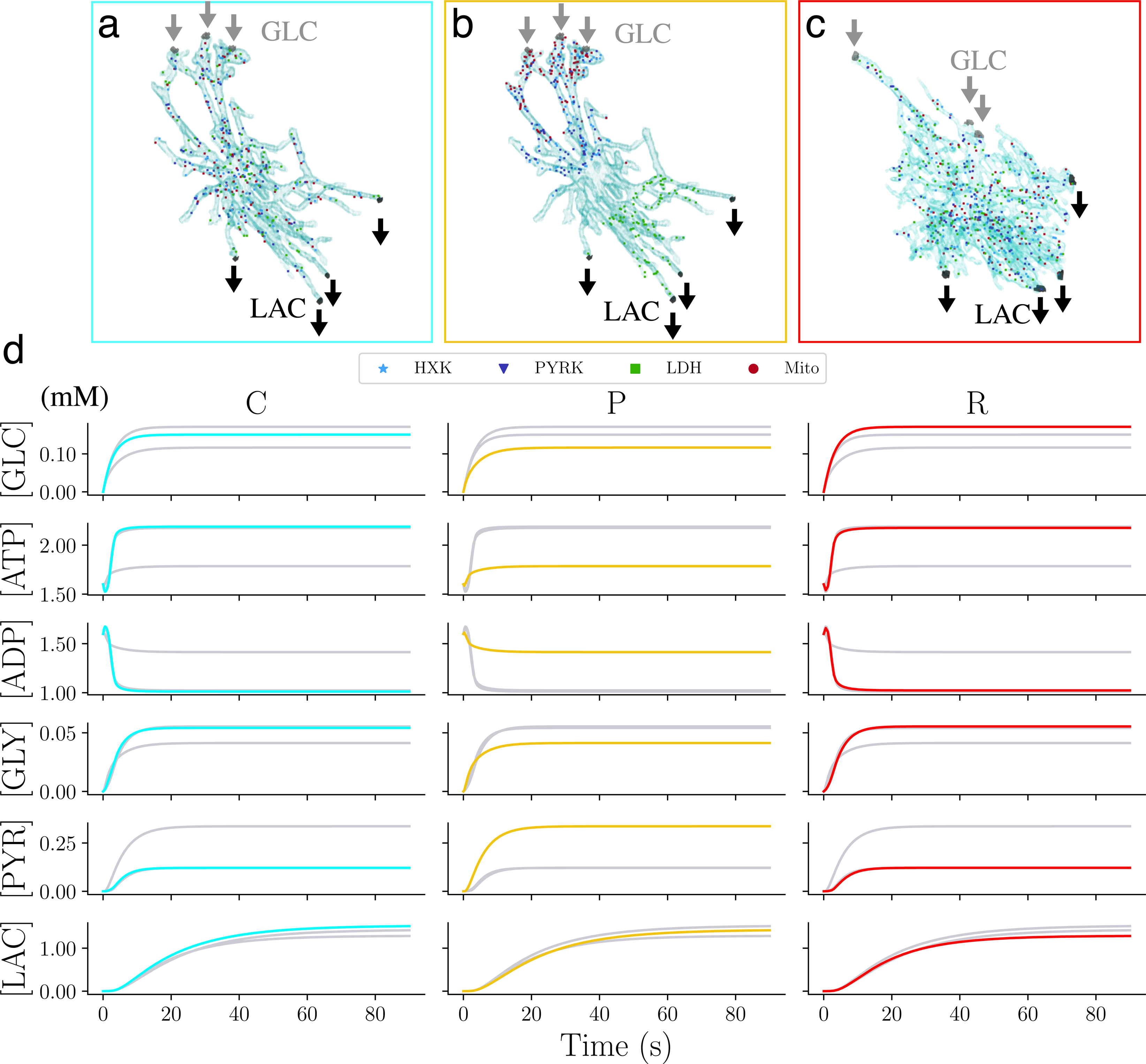
Metabolite dynamics in 3D astrocytes with physiological reaction site versus extreme polarised arrangements. Setting of the 3D simulations for the **a** control (C), **b** polarised (P) and **c** reactive (R) astrocyte. For C and R, Mito reaction centers were inferred from the microscopic images. Each HXK site is sorted from a gaussian distribution centered at each Mito site. In this way for each Mito we have an HXK reaction close by. PYRK and LDH are uniformly distributed inside the box that contains the cells. The reaction centers of P are sorted in the way that HXK and PYRK are colocalised close to the GLC influx, while on the other extremity of the cell we locate LDH centers. Mito centers are sorted using a log-normal distribution that locate them in the same region as HXK. The number of centers per reaction type is 90 for C and P, and 140 for R. For the three settings, GLC enters three sub-regions from the branches of the cell in contact with the blood vessels and LAC exits from four sub-regions at the other extremity of the cell. **d** Time behavior of the average concentration of each species for C (cyan), P (yellow) and R (red).

To investigate a reactive astrocyte subject to AD, we extended our simulations by gradually adding AD-related dysfunctions. Experiment 1 (E1) mimics a loss of GLC uptake ^31^ by a 30% decreased GLC influx. Experiment 2 (E2) includes the dysfunction in mitochondrial activity ^32–34^ inducing a lower ATP production by a reduced reaction rate for Mito (*K*_Mito_10*^−^*^5^). In accordance with available experimental data ^8^, we considered an increment of the activity of LDH by a factor of ten in Experiment 3 (E3) and an increment in the glycolysis rate in particular in the PYRK reaction, also by a factor of ten in Experiment 4 (E4). In the final experiment (EAD), all four conditions were combined to explore their possible synergistic effects.

Fig. 7**a** exhibits the percentage of the concentration loss at steady state for experiments E1, E2, E3, E4 and EAD compared to R. Interestingly, the 30% reduction in GLC uptake in E1 is reflected by the final steady state in GLC (*≈* 28.7% loss) which induced a loss of *≈* 35% in GLY, PYR and LAC. Dysfunctional Mito reactions lead to an increase in final GLC level and a loss in ATP and GLY whereas the level of LAC is not affected. The experimentally observed increased activity of LDH (considered in experiment E3) results mainly in faster metabolizing of PYR. On the other hand, GLY consumption is maximised by the turnover of PYRK in the E4 experiment while the other concentrations are not affected. The combined effect of the individual dysfunctions in the EAD experiment leads to a significant change in the metabolic profile (with the highest loss in ATP, GLY and PYR Fig. 7**a**). (The dynamics of these experiments is shown in Supplementary Note 4.)

**Fig.7.**
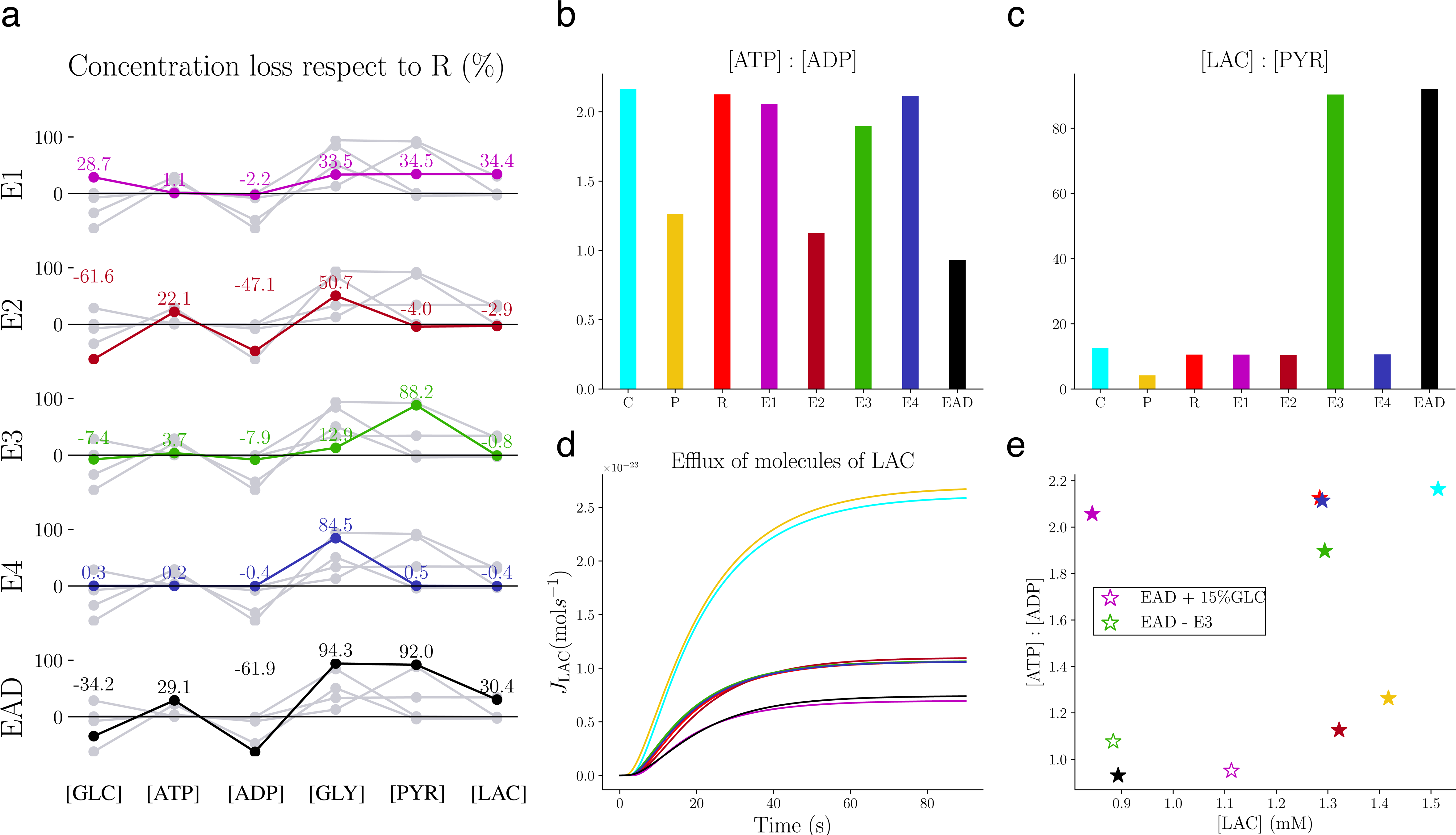
Effects of AD conditions on metabolic average concentrations. We consider four pathological conditions of AD, in the setting of the reactive astrocyte R. E1 describes the deficiency of GLC uptake (magenta); E2, the mitochondria dysfunction (dark red); E3, the LDH overwork (green); E4, PYRK overwork (blue) and EAD, the four conditions combined (black). **a** Final average concentration loss respect to R. The experiments reflect their loss/gain imposed to the cell through the conditions. Steady-state spatially averaged **b** ATP : ADP ratio and **c** LAC : PYR ratio of control (cyan), polarised (yellow), reactive (red) and all the AD experiments. **d** Efflux of LAC molecules exported over time from the astrocyte to the extracellular space. The experiments with higher export are the two control astrocyte with C and P configurations. The experiments with a lower export are E1 with a loss in GLC uptake and EAD with the combination of the AD conditions. **e** Scatter plot of ATP : ADP against LAC final average values. The most efficient cell is the control one. Then, the different AD conditions affect the cell status leading the reactive cell affected by all the AD conditions to an unhealthy state. In order to save the EAD cell, we increase the uptake of GLC up to 85% (white star with magenta edge), and the cell responds by using the more available fuel to produce more LAC. However, blocking the LDH overwork (white star with green edge) increases ATP : ADP and thereby rescues the astrocyte from the AD conditions.

The functional state of cells in terms of ATP : ADP and LAC : PYR ratios at steady state is preserved for a wide range of conditions. Even for the polarised P configuration, the ATP : ADP ratio is higher than 1.0 (Figs. 7**b** and 7**c**), suggesting that a complex shape makes the cell more robust against extreme situations. This is also confirmed by the ratios of the E2 experiment, that does not exhibit a ratio below 1.0 despite mitochondrial dysfunction. The only cell that reaches a critical unhealthy state is the EAD condition (0.93), where mitochondrial dysfunction adds to the other dysfunctions. Also the ratio of LAC : PYR is always within physiological range (*>* 10) for all conditions except P. However, a LAC : PYR ratio of above 80 are reported in E3 and EAD, which may indicate hypoxia with low levels of oxygen in blood ^16^.

**Fig.8.**
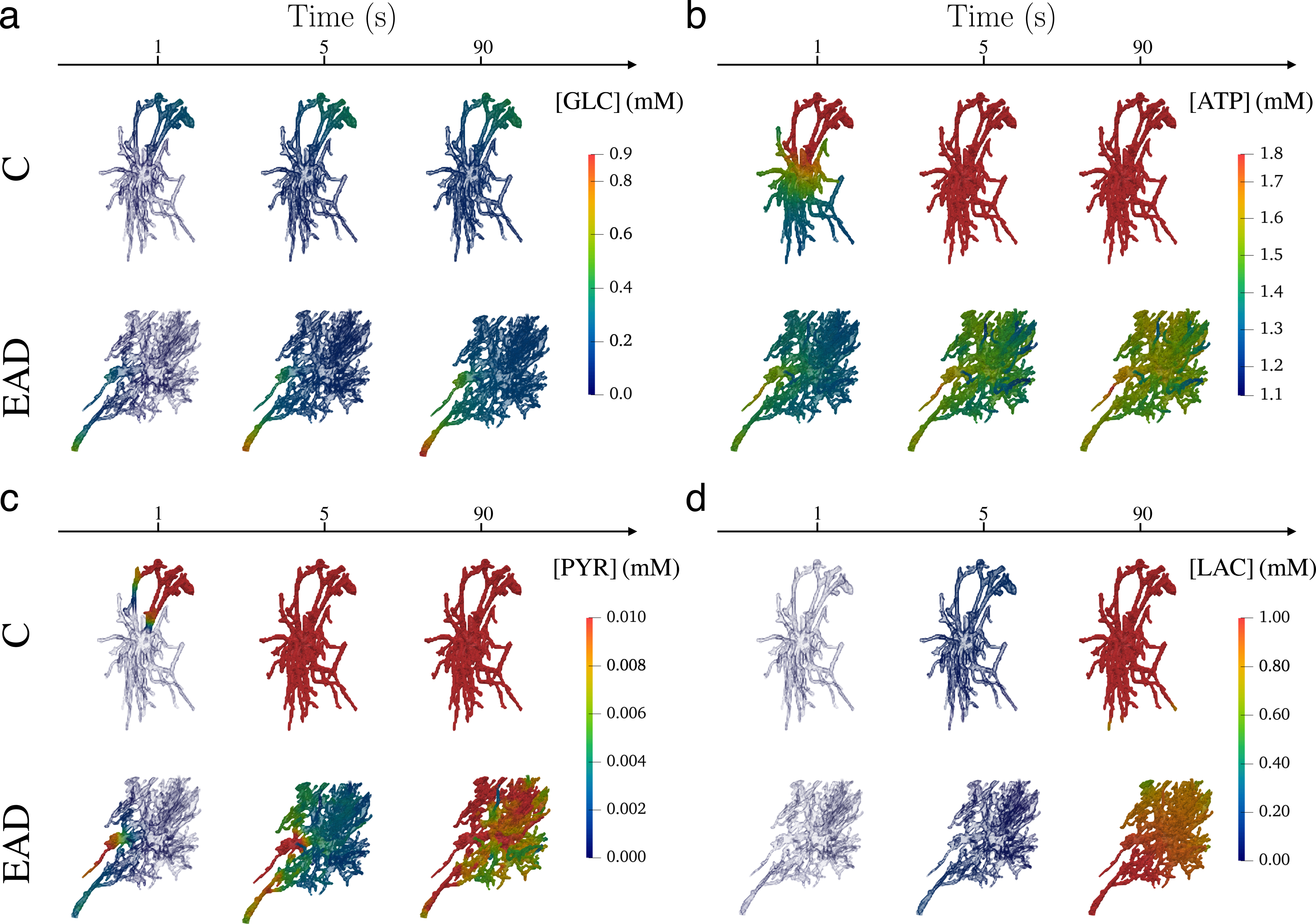
Spatially resolved Control and EAD astrocytes for GLC, ATP, PYR and LAC at different times. 3D spatial concentration of metabolites at three different time steps in control (C) and reactive astrocyte affected by AD pathology (EAD). **a** GLC enters from the blood vessels and spreads inside the astrocytic domains activating the glucose metabolism. **B** ATP, already present in the cells at the initial time, is produced and consumed. In particular, in correspondence with regions with high numbers/absence of Mito sites, we can notice high/low levels of ATP in EAD. **c** PYR produced by PYRK diffuses inside the 3D domains and highlight the complex shape of the reactive astrocyte with high variability of concentration within the cell. **d** LAC shows a slow production, in fact at time 5, both C and EAD show low concentrations. At the final time, we can appreciate the steady state level of LAC where the regions where it is exported are highlighted by lower concentrations.

Since LAC export into the extracellular space is an essential mechanism of astrocytic support to neurons, we also quantified LAC efflux exporting LAC from the corresponding subregions (Fig. 7**d**). The asymptotic behaviour of the efflux indicates that cells with the C and P configuration export more LAC, suggesting that the less ramified morphology of the protoplasmic astrocyte allows for faster diffusion of molecules and subsequent export regions. On the other hand, E1 and EAD configurations export less, indicating that the 70% decrease in GLC uptake might drive this AD symptom. The different metabolic states of the cell are also assembled in the “altruistic” vs “egocentric” map in terms of the LAC concentration and the ATP : ADP ratio (Fig. 7**e**). This map indicates the C configuration as the most efficient cell with high levels for both in agreement with the previous finding on uniform distributed cells. The P setup exhibits a more altruistic behaviour than expected by producing more LAC than ATP, potentially facilitated by the morphology. When cells lack GLC, they become more egoistic and produce more ATP. Remarkably, the steady state of LAC of the R, E2, E3 and E4 experiments is *≈* 1.3 *µ*M but the ATP concentration is decreasing from high levels in the R and E4 configuration to lower concentration in E2. Finally, lower levels of both ATP and LAC is the AD-related EAD condition suggesting that it can neither support neurons nor itself. Last, we studied how to support an AD-impacted astrocyte where the results of the individual conditions helped to disentangle the different effects. Importing more GLC (by increasing the uptake to 85% of the healthy control condition) turns the cell into a more altruistic state by using the additional fuel predominantly for LAC production. Blocking the excessive activity of LDH saves the cell from AD-related energy deprivation but with the cost of reduced LAC export.

To investigate the impact of diffusion limitation as an underlying mechanism in reactive astrocytes, Fig. 8 illustrates the time evolution of the 3D distribution of concentrations for the healthy C and AD-related EAD condition considering the properties and spatial distribution of reaction sites (Supplementary Movies 1-12). In particular, the trapping effect discussed above is highlighted in the reactive astrocytic morphology for ATP and PYR where branches exhibit a higher concentration variability.

To summarise, the physiologically realistic simulations reproduce important features of astrocytes in healthy and diseased conditions. The incorporation of real morphologies highlights cellular robustness against extreme enzymatic configurations. This is also seen for AD conditions, indicating the influence of the cellular domain on the metabolic state of the cell. In fact, a single AD characteristic does not lead to an unhealthy cell, only combinations of AD-terrain leads to severe metabolic dysfunctions.

## Discussion

Although the link between cellular morphology and metabolic activity might have implications for neurodegeneration including Parkinson’s disease and Alzheimer’s disease, our understanding of this connection remain imperfect due in part to experimental limitations. To address this challenge, we developed a multiscale model for energy metabolism in complex cellular domains with a specific focus on the intracellular spatial orchestration of astrocytes. To build the mathematical model, we first considered a single reaction site for each metabolic subpathway in a 2D circular geometry and validated the model in terms of physiological concentration ranges for astrocytes (Fig. 1), in accordance with previous ODEs model ^20^. We showed numerically that different spatial organisation of reaction sites lead to distinct metabolic profiles due to diffusion limitation and local substrate competition (Fig. 2). The observed differences between the circular and the star-shaped domain indicated a possible trapping effect for molecules in more complex shapes. These trapping effects might be overestimated compared to a more physiologically realistic astrocyte, since many more reaction sites are typically present within an astrocytic branch. Nevertheless, these results strongly indicate that the spatial dimension and the domain complexity can have a crucial effect on metabolic profiles and may be of particular importance for the metabolic support function of astrocytes.

To further characterize these spatial effects in a more physiological setting, we considered a larger number of reaction sites, which were distributed either within a uniform or polarised arrangement inside a rectangular shape, mimicking an astrocytic branch. For each configuration, we ran 200 realisations, allowing for robust statistical comparisons between the different settings (Fig. 3). The results showed that cells with uniformly distributed reaction sites are significantly more efficient in both the “altruistic” LAC production as well as the “egocentric” intracellular energy state. Although polarised organization corresponds to an extreme and rare biological setting, the analysis of these realisations indicates the importance of a more homogeneous mitochondria distribution for a sufficient activity and a related energized cell state (Fig. 4). This is in line with experimental observation of mitochondrial organisation and homeostasis including fission and fusion where impairment of these processes are linked to neurodegeneration ^35^.

Based on the 2D model, we extended our investigations to physiological 3D morphologies of astrocytes, obtained from confocal microscopy images of human *post-mortem* brain samples of an AD patient and a healthy control subject. Our approach is thereby able to integrate directly the spatial orchestration of reaction enzymes as demonstrated by the experimentally quantified mitochondrial distribution (Fig. 5). We first confirmed that using different morphologies but the same parameters lead to concentrations in the physiological range in agreement with the findings in the simplified 2D geometries (Fig. 6). To investigate the effect of AD-related molecular modifications, we analysed a reactive astrocyte with baseline parameters and four individual metabolic dysfunctions linked to AD and their combinations (Fig. 7). The results highlighted that different pathological effects arouse specific system response and differentiated the cell behaviour between an “altruistic” and an “egocentric” mode. Furthermore, the results indicated that any give dysfunction does not lead necessarily to a dysfunctional cell with a low ATP : ADP ratio but it is the cumulative metabolic insufficiencies that lead the cell into a critical state. This synergistic phenotype might be related to the multi-hit perspective in neurodegeneration which addresses the transient compensation and typical disease onset at higher age ^36, 37^. The systematic study of the individual dysfunctions allowed to suggest that reducing LDH activity could sustain astrocytic function. Such approaches are also discussed in the context of cancer ^38, 39^. However, in the context of AD, the challenge would be to interfere with metabolism in a cell-type specific manner.

Furthermore, the comparison between the simplified 2D domains and the complex 3D morphologies indicates that real astrocytic shape affect the cell state with robustness towards enzyme orchestration and different metabolic dysfunctions. This robustness might be caused by the trapping of molecules in thin branches as further indicated by the analysis of 2D star-shaped morphology. The thin processes may hamper the diffusion of molecules as shown by the spatial concentra-tion profiles (Fig. 8) which increased mitochondrial activity and corresponding ATP production with the cost of decreased LAC export. Thus, the complex morphology might provide a mechanism to support an “egocentric” state if the system reaches limiting conditions, similar to energy buffering in complex mitochondrial morphologies ^40^.

To our knowledge, our approach is the first 3D model of cellular energy metabolism using physiological human cellular morphologies. Our analyses of hippocampal control and AD-related reactive astrocytes clearly demonstrate the importance of morphology for cellular metabolic activity. Our approach has limitations, such as the lack of cellular compartmentalisation, the coarsegraining of enzymatic reaction into effective metabolic pathways, the limitation of the GFAP staining and the incomplete information on reaction site localization provided by imaging modalities. Despite these limitations, we demonstrate the general importance and feasibility of physiological simulations by integrating molecular properties, spatial intracellular orchestration and morphology. Based on our multiscale framework, future investigations will allow to disentangle different mechanisms underlying neurodegeneration, including mitochondrial morphology ^40, 41^, organization and dysfunction ^42–44^ by more detailed models.

## Methods

### Energy Metabolism Model

The core energy metabolism is broken down into the core metabolic pathways by the coarse-grained non-reversible reactions:

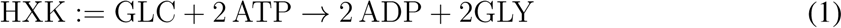

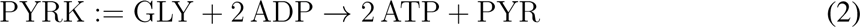

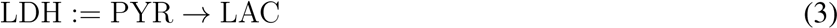

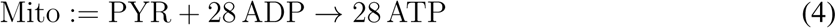

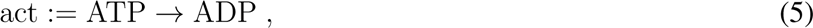

where the first two reactions consider the ATP consuming and ATP producing parts of glycolysis, LDH describes the activity of lactate dehydrogenase. Mito reflects the overall metabolic activity of mitochondria in terms of ATP production and general cellular activity is reflected by the act reaction.

### Reaction Diffusion System

To investigate the spatial coupling of the metabolic pathways (Eqs. (1)- (5)), the reactions were integrated by a RDS ^45^. The domain of the PDEs is a bounded subset of R*^d^* (*d* = 2 or 3), denoted by Ω and concentrations [ *·* ] are defined as function [ *·* ] : Ω *×* [0*, T* ] *→* R. Diffusion coefficients for each species are given by *D*_[_ *_·_* _]_ and chemical reactions are modeled using mass action kinetics ^46^. The reaction rate for homogeneous cellular activity (*K*_act_) and a spatial reaction rate density, *K_j_*, for the other four reactions. Considering *M* reaction sites located in *{***x***_i_}^M^ ∈* Ω, the spatial reaction rates are defined as the product between the classical reaction rates, *K_j_*, and Gaussian functions located at those reaction sites with variance *σ_i_ ∈* R^+^:

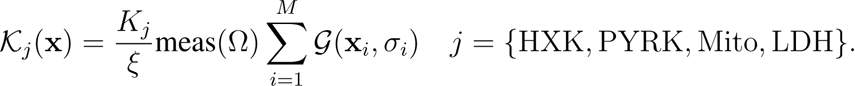

*ξ* is a parameter that ensure the property that ^J^_Ω_ *K_j_*d*x* = *K_j_* and meas(Ω) is the area of the domain in 2D or the volume in 3D. The source of GLC is described through a function *J*_in_ : Ω *×* [0*, T* ] *→* R:

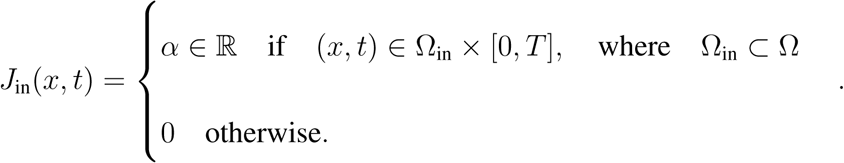

Similarly, the degradation of LAC, which is proportional to the amount of LAC in region Ω_out_ *⊂* Ω is described by function *η*_LAC_ : Ω *×* [0*, T* ] *→* R

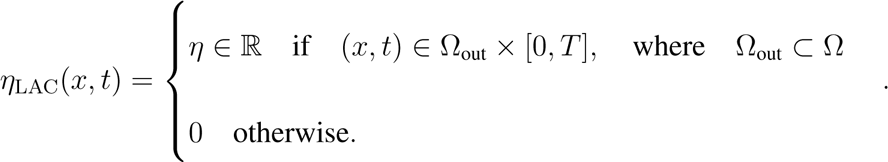

With this definition the reaction diffusion system is given by

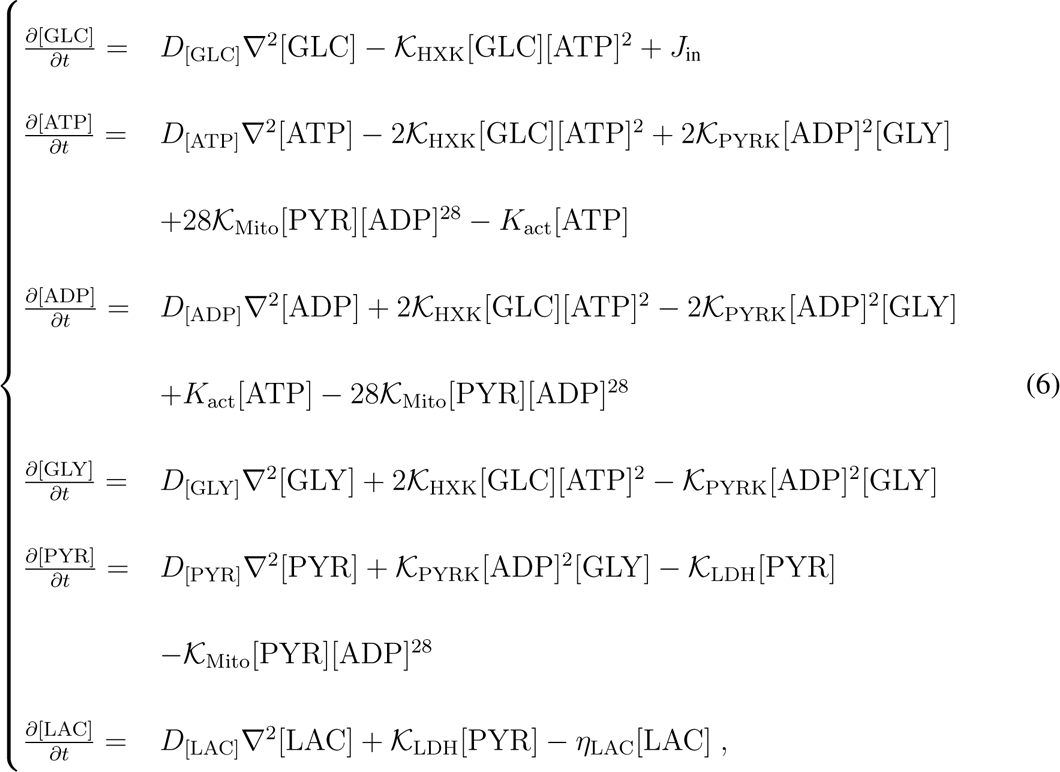

where we considered von Neumann boundary condition to consider no-flux settings at the cell membrane. To characterize the system’s behavior, we analysed the equilibrating dynamics towards the steady state from the initial conditions for ATP and ADP concentrations

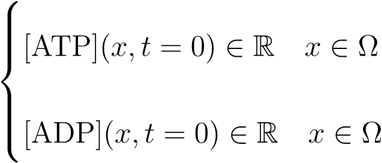

where an initial ATP concentration is required for the initial glycolysis reactions and vanishing concentrations for the other species. To ensure robust simulations, we transformed the RDS into a dimensionless system allowing for convergence over a large parameter range (Supplementary Note 5).

### Human brain tissue

Post-mortem brain tissue was obtained from the Douglas-Bell Canada Brain Bank and handled according to the agreements with the Ethics Board of the Douglas-Bell Brain Bank (Douglas Mental Health University Institute, Montréal, QC, Canada) and the Ethic Panel of the University of Luxembourg (ERP 16-037 and 21-009). The two hippocampal samples used in this work were donated from a male 87-year-old Alzheimer’s Disease patient with a disease stage of A2B3C2 and a post-mortem interval of 21,75 hours, and by a female 89-year-old (age-matched) control subject with a post-mortem interval of 23,58 hours.

### Immunofluorescence stainings

The PFA-fixed hippocampal samples were cryosectioned into 50 *−* 100 *µ*m thick slices on a sliding freezing microtome (Leica SM2010R). To visualize astrocytes and mitochondria, we co-immunostained the slices against glial fibrillary acidic protein (GFAP) and Tu translation elongation factor mitochondrial (TUFM) respectively. The target-binding primary antibodies used here were Anti-GFAP guinea-pig (Synaptic Systems Cat# 173 004, RRID:AB 10641162) at a dilution of 1:500, and Anti-TUFM mouse (Atlas Antibodies Cat# AMAb90966, RRID:AB 2665738) at a dilution of 1 : 200. The corresponding fluorophore-coupled secondary antibodies used were Alexa Fluor 647-AffiniPure Donkey Anti-Guinea Pig IgG (H+L) (Jackson ImmunoResearch Labs Cat# 706-605-148, RRID:AB 2340476) at a dilution of 1 : 300 and Alexa Fluor 488-AffiniPure Donkey Anti-Mouse IgG (H+L) (Jackson ImmunoResearch Labs Cat# 715-545-150, RRID:AB 2340846) at a dilution of 1 : 400. We followed a previously published protocol ^24^ with the exception of a double incubation with primary antibodies for the TUFM staining.

### Image acquisitions

High-resolution confocal images with 0.333 *µ*m z-step were acquired using a Leica DMi8 microscope with a 93X glycerol objective and LAS X software (Leica Microsystems). The region of interest was fixed on the hippocampal subregion CA1.

### Image pre-processing

The surface function of Imaris 9.6.0 software was used to segment GFAP staining to produce astrocyte morphology 3D reconstructions. The surface grain size parameter was set to 0.3 *µ*m for the segmentation of astrocyte morphology. Upon segmentation of the GFAP signal of the entire image, we manually selected the astrocyte of interest and removed all other non-relevant segmentation structures. The spots function was used to segment TUFM staining. The estimated spots diameter was set to 0.2 *µ*m. To select only the mitochondria of interest (corresponding to the astrocyte of interest) we applied the filter of the spots function called ‘Shortest Distance to Surface’ [segmented astrocyte]. In the control astrocyte some mitochondria of interest were not automatically selected by this filter setting, because they were too far away from the segmented surface, however part of the astrocyte, notably in the cell soma. To include these mitochondria into the analysis, a second filter was applied twice by selecting the central mitochondria of the soma compartment and applying ‘Shortest Distance to Surface’ function.

The direct use of the astrocytic segmented images as domain for our simulation would require a mesh fine enough to capture the thin branches of the cellular structure. This would mean billions of quality finite elements, with a good aspect ratio. In literature, this problem was addressed by refining the mesh in critical regions ^47^. However in our case, this would require refining all branches. We overcome these issues by additional image pre-processing where we dilated and down-sampled the binary images. These two steps enlarged the thin branches and avoid discontinuities when we map the images to the finite element mesh. These steps are not critically affecting the real morphology of the astrocytes and might actually address partially the GFAP staining limitation. Moreover, we impose the astrocytic volume in the simulations to be equal to the one of the segmented images obtained with Imaris. Eventually, we obtained the final segmented images (*f* ) with labeling the voxels inside (*−*1), outside (1) and on the boundary (0) of the astrocytes.

Before applying the same steps to the binarised segmented mitochondrial images, we applied a convolution filter to smooth the voxels. To extrapolate the information about mitochondrial density, we selected all connected components in the images and for each of them, we identified the center and the radius of the circle that contains such component.

### Numerical methods

To solve numerically the RDS, the first step is to convert Eq. 6 into a corresponding weak form ^48^. Then, we discretise the weak form both in time and space. we discretise the time derivative using a finite difference method (backward Euler) ^49^ and the spatial domain by finite elements ^50^ and cut finite elements ^51^. The 2D experiments were solved using classical finite element methods based on FENICS ^52, 53^, while the 3D experiments were solved using CUTFEM ^26, 51^. Since the weak RDS formulation is non-linear, we linearised it and used a Newton-Raphson algorithm to iteratively solve the problem. The linear system at each time step of the Newton-Raphson algorithm was solved using standard linear solver from the *PETSc* library. For further details and numerical parameters see Supplementary Notes 6 and 7, respectively.

### Physiological model parameters

The parameters used in our model are given in Table 1. The diffusion parameters were chosen for ATP and ADP following ^54^, for GLC based on ^55^ and for the other species based on the Polson method ^56, 57^.

The calibration of the reaction rates has been done in accordance with the steady states of the ODE system ^20^ associated to Eq. (6). For *J*_in_ we used the maximum transport rate of GLC from ^20^. For *J*_out_ we used the maximum transport rate of LAC but divided it for the steady state, since we required our transport of LAC to be proportional to the local concentration of LAC inside the cell.

## Acknowledgement

Authors would like to thank Corrado Ameli for his help in the segmentation of the control astrocyte and Mark Ellisman (UCSD) and Jack S. Hale for fruitful discussions. S Farina and S Fixemer were supported by the PRIDE program of the Luxembourg National Research Found through the grants PRIDE17/12252781/DRIVEN and PRIDE17/12244779/PARK-QC, respectively.

## Contributions

AS and SB conceptualized the research. S Farina contributed to the software, formal analysis and visualization of the results. S Farina, VV, SB and AS contributed to investigation of the results. S Fixemer and DB provided biological resources and contributed to the visualization. SC contributed to the CUTFEM methodology. S Farina, S Fixemer and VV wrote the initial draft. All authors contributed to reviewing and editing the final manuscript. AS and SB supervised the project.

