## Supplementary Information for "Mechanistic multiscale modelling of energy metabolism in human astrocytes indicates morphological effects in Alzheimer’s Disease"

#### Supplementary Note 1: Spatial arrangements for 2D simulations in rectangular shape.

In the rectangular domain we set up our simulation with ten reaction sites per reaction type. We define the entrance of $\mathrm{GLC}$ at the origin, the bottom left corner and the exits of $\mathrm{LAC}$ on the opposite vertex $(4, 140)\,\mu\mathrm{m}$. The subregions of entrance and exits have been defined as the intersection of the rectangle and the circle with center the origin or the top right corner and radius $1.0\,\mu\mathrm{m}$.

We present in Supplementary Table 1, the different distributions used to define the $x$ and $y$ coordinates of the enzyme arrangements inside the 2D rectangle ($[0, l_1] \times [0, l_2]$). As presented in the main text, each setting has 10 reaction sites per reaction type. The uniform cells have all their sites sorted from a uniform distribution, noted $\mathcal{U}[a, b]$ covering the whole rectangular domain.

In polarised cells, we assumed the enzymes to be distributed according to normal distribution ($\mathcal{N}(m, \sigma')$, where $m$ and $\sigma'$ are the mean and standard deviation, respectively) or log-normal ($\log \mathcal{N}(m, \sigma')$) distributions. Close to the $\mathrm{GLC}$ influx we place $\mathrm{HXK}$ and $\mathrm{PYRK}$ reactions, using $\mathcal{N}(\frac{l_1}{2}, 2)$ and $\mathcal{N}(10, 5)$ to select the $x$ and $y$ coordinates, respectively. The ten enzymes of $\mathrm{LDH}$ are located close to the $\mathrm{LAC}$ efflux with $(x, y) \in (\mathcal{N}(\frac{l_1}{2}, 2)), \mathcal{N}(l_2 - 10, 5)$. For the ten mitochondria, we select the $x$ coordinate as for the other reaction sites with a normal distribution $\mathcal{N}(\frac{l_1}{2}, 2)$. For polarized cells, one option consists in locating six reacting sites with $y$ selected from a normal law $\mathcal{N}(10, 5)$ and distributing the remaining four uniformly within the top part of the cell ($y > 60$). We refer to this setting as the "Polarised" one. We also consider a different sampling where mitochondrial locations are distributed according to a log-normal law $\log \mathcal{N}(2., 2)$. We call this setting "Polarised $\log \mathcal{N}(2)$".

Typical polarised configurations are although lacking mitochondria in the middle part of the cell as shown in the corresponding figure of the main text.

| Reaction site distributions | | | | | | |
|---|---|---|---|---|---|---|
| | Uniform | | Polarised | | Polarised $\log \mathcal{N}(2)$ | |
| | $x$ | $y$ | $x$ | $y$ | $x$ | $y$ |
| HXK | $\mathcal{U}_{[0,l_1]}$ | $\mathcal{U}_{[0,l_2]}$ | $\mathcal{N}(\frac{l_1}{2}, 2)$ | $\mathcal{N}(10, 5)$ | $\mathcal{N}(\frac{l_1}{2}, 2)$ | $\mathcal{N}(10, 5)$ |
| PYRK | $\mathcal{U}_{[0,l_1]}$ | $\mathcal{U}_{[0,l_2]}$ | $\mathcal{N}(\frac{l_1}{2}, 2)$ | $\mathcal{N}(10, 5)$ | $\mathcal{N}(\frac{l_1}{2}, 2)$ | $\mathcal{N}(10, 5)$ |
| LDH | $\mathcal{U}_{[0,l_1]}$ | $\mathcal{U}_{[0,l_2]}$ | $\mathcal{N}(\frac{l_1}{2}, 2)$ | $\mathcal{N}(l_2 - 10, 5)$ | $\mathcal{N}(\frac{l_1}{2}, 2)$ | $\mathcal{N}(l_2 - 10, 5)$ |
| Mito | $\mathcal{U}_{[0,l_1]}$ | $\mathcal{U}_{[0,l_2]}$ | $\mathcal{N}(\frac{l_1}{2}, 2)$ | $\mathcal{N}(10, 5)$ or $\mathcal{U}[60, l_2]^*$ | $\mathcal{N}(\frac{l_1}{2}, 2)$ | $\log \mathcal{N}(2, 2)$ |

Table 1: The table shows the distributions chosen for $x$ and $y$ coordinates for each metabolites for the simulations in the 2D rectangle for the Uniform, Polarised and Polarised $\log \mathcal{N}(2)$ cells. * indicates that we sorted the $\mathrm{Mito}$ sites from two distributions: six from the normal and four from the uniform. In this way, we ensure the probability of having four $\mathrm{Mito}$ sites on the top of the rectangle.

**Supplementary Note 2: Significance test for 2D realisation**

We run 200 realisations for each of these configurations (*i.e.* uniform, polarised and polarised $\log \mathcal{N}(2)$ for the purpose of statistical testing. In order to find out if the three types of spatial arrangements lead to statistically significant differences in the realisations, we perform a multiple comparison Holm-Bonferroni method since we consider simultaneously the distribution of the different concentrations. The Bonferroni method is applied to a parametric independent T-test to evaluate if there is a significant distance between the means of the concentrations of the three configurations and to a non-parametric Wilcoxon-Mann-Whitney test to verify if two statistical samples come from the same population. The results of the significance tests were presented in Table 2. When the $p-$value is smaller than $0.05$ then the hypothesis of the tests is rejected, meaning that our samples describe different populations, which is mainly the case.

The p-value results show that only $\mathrm{GLY}$ for the uniform and polarised cell, and $\mathrm{PYR}$ for the two polarised configurations are not significantly impacted by the spatial arrangements where $\mathrm{GLY}$ exhibits a high $p-$value only in the t-test but not in the non-parametric one. This finding is consistent with the similar average of the steady state concentration but the distinct underlying distribution (Fig. 3**c**). For $\mathrm{PYR}$, the distributions for the two polarised cells exhibit a similar range (Fig. 3**c**) but since the two polarised cells only differ by the distribution of mitochondria it is expected that the distribution of $\mathrm{PYR}$ is similar, since it is produced in $\mathrm{PYRK}$ reaction.

**Supplementary Note 3: Spatial arrangement for 3D simulations.**

We present in detail the settings of the simulation shown in Figs. 6**a**-**c**. For the control, C, and the reactive, R, $\mathrm{LDH}$ and $\mathrm{PYRK}$ sites were sorted by a uniform distribution defined in the boxes that contains the two astrocytes, see Fig. 1. While $\mathrm{HXK}$ sites were sorted from a normal distribution centered in each $\mathrm{Mito}$ site and with variance $0.03 \times L$ where $L$ is a dimensionless parameter (see below Supplementary Note 5). In particular, $1.38\,\mu\mathrm{m}$ for the control and $2.75\,\mu\mathrm{m}$ for R.

The polarised settings were arranged colocalising $\mathrm{HXK}$ and $\mathrm{PYRK}$ enzymes from a uniform distribution that cover the endfeet of the astrocyte ($\mathcal{U}_{[0,l_x]\times[y_1,l_y]\times[0,l_z]}$, where $l_x, l_y, l_z$ are the dimensions of the box that contains the control astrocyte). $\mathrm{LDH}$ were sorted from the opposite side of the box containing the $\mathrm{LAC}$ export sub-regions of the astrocyte from another uniform distribution $\mathcal{U}_{[0,l_x]\times[0.,y_2]\times[0,l_z]}$. While the $\mathrm{Mito}$ were sorted from a log-normal distribution $\log\mathcal{N}(l_x, 0.64, l_z, 0.2)$ in the manner that colocalise $\mathrm{HXK}$ and $\mathrm{PYRK}$.

Last, the three sub-regions chosen for the $\mathrm{GLC}$ entrance and the four $\mathrm{LAC}$ exits were selected manually and they are defined as the intersection of the cellular morphology with a sphere with a radius of $1.0\,\mu\mathrm{m}$.

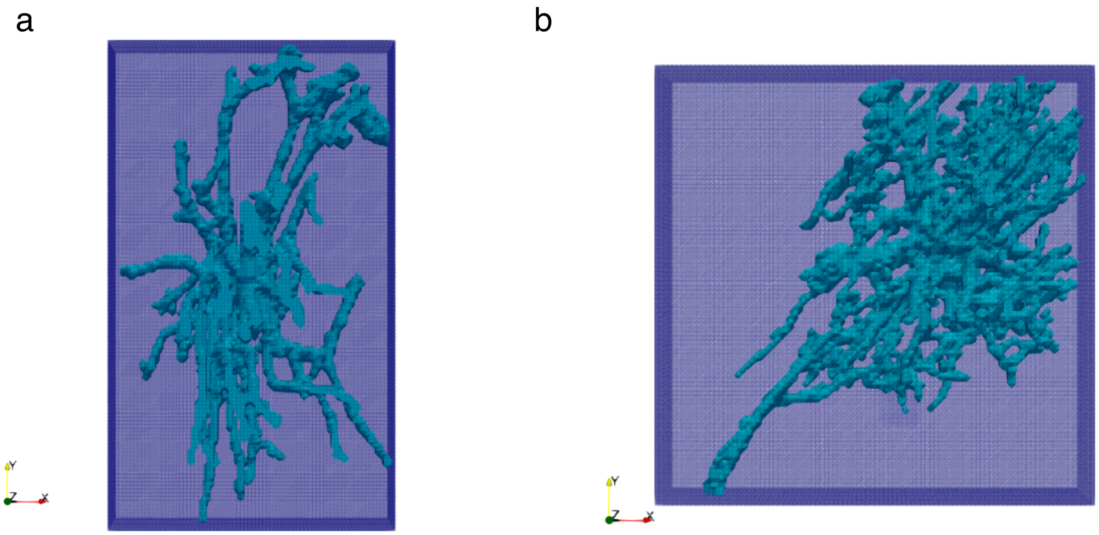
a
Y
Z
X
b
Y
Z
X

**Figure 1 Supplementary Fig.1: Astrocytic morphologies embedded into a finite element background mesh. a** control and **b** reactive AD astrocytes are implicitly defined using a level set function and embedded in a structured background mesh. This is how we separated the finite element mesh from the geometries of the objects. Moreover, we used the dimensions of the background meshes to define the bounds of the uniform distributions to sort some of the reaction site centers.

### Supplementary Note 4: Additional Figure for AD simulations

We present in Supplementary Fig. 2 an extension of Figure 7, where we show the trajectories of the average concentrations of the metabolites. This plot highlights the behaviour of the system in response to different AD related conditions.The systems reach the steady state in $\approx 50(\mathrm{s})$.

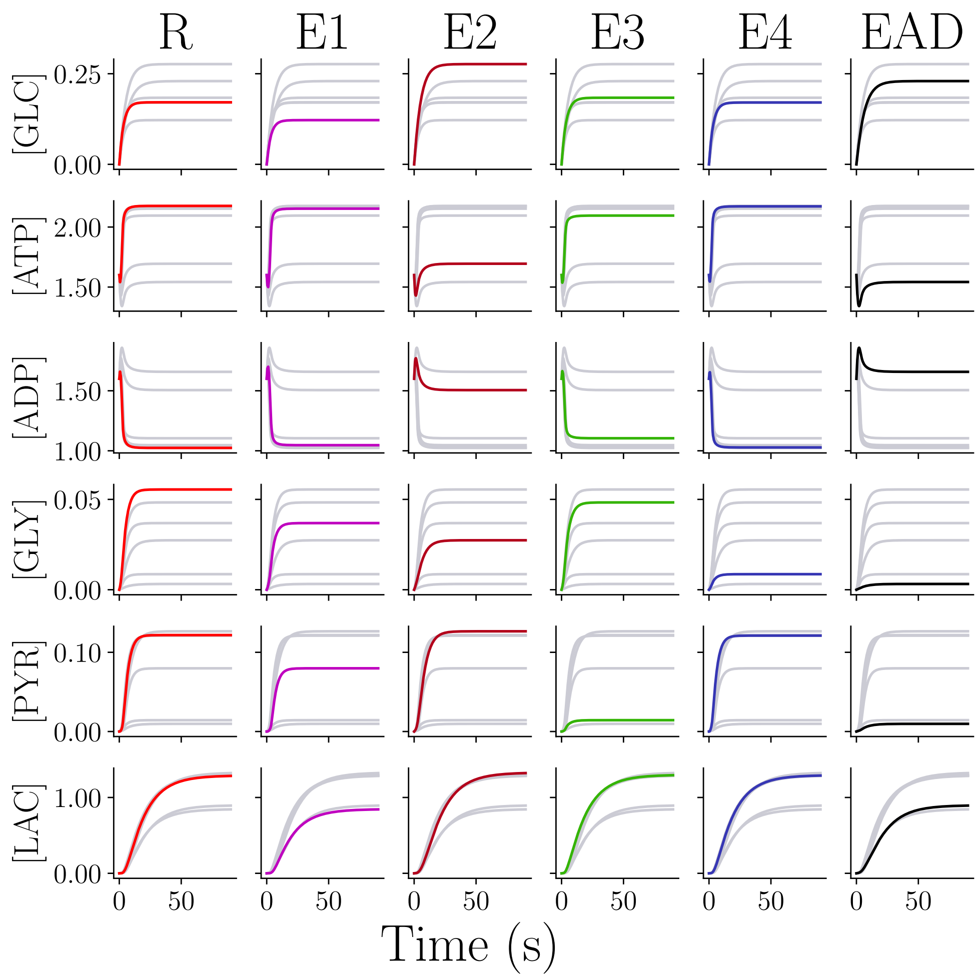


**Figure 2 Supplementary Fig.2: Effects of AD conditions on metabolite dynamics in 3D reactive astrocyte.** Dynamics of the average concentration of each metabolites the simulations are solved inside the reactive astrocyte in AD with the setting presented in Figure 6 **c** of the main manuscript. R is the solution obtained with healthy parameters presented in Table 1 (red). E1 describes the deficiency of $\mathrm{GLC}$ uptake (magenta); E2, the mitochondria dysfunction (dark red); E3, the $\mathrm{LDH}$ overwork (green); E4, $\mathrm{PYRK}$ overwork (blue) and EAD, the four conditions combined (black).

**Supplementary Note 5: Dimensionless system.**

To obtain the dimensionless system from the RDS Eq. 6, we impose $[\bar{\text{GLC}}] = \frac{[\text{GLC}]}{\alpha}$, $[\bar{\text{ATP}}] = \frac{[\text{ATP}]}{A_{tot}}$, $[\bar{\text{ADP}}] = \frac{[\text{ADP}]}{A_{tot}}$, $[\bar{\text{GLY}}] = \frac{[\text{GLY}]}{\alpha}$, $[\bar{\text{PYR}}] = \frac{[\text{PYR}]}{\alpha}$, $[\bar{\text{LAC}}] = \frac{[\text{LAC}]}{\alpha}$, where $A_{tot}$ is the total amount of ATP and ADP inside the cell and $\alpha$ a constant that we set to $0.16$. The spatial dimensionless is $\bar{x} = \frac{x}{L}$ $\bar{y} = \frac{y}{L}$ and $\bar{z} = \frac{z}{L}$ where $L$ is a parameter based on the volume of the astrocyte:

$$L = \frac{V}{V_\Omega}$$

where $V$ is the real volume of the astrocyte segmented and $V_\Omega$ is the volume of the astrocyte in the dimensionless box with fixed $x$ size of $1.0$. We also define the dimensionless time parameter as $\bar{t} = \frac{t}{t_c}$ and we choose $t_c = \frac{L^2}{D_{\text{GLC}}}$. So, we have that the dimensionless system is:

$$
\begin{cases}
\frac{\partial[\bar{\mathrm{GLC}}]}{\partial\bar{t}} = & \nabla^2[\bar{\mathrm{GLC}}] - \mathcal{K}_{\mathrm{HXK}}\beta_{[\bar{\mathrm{GLC}}]}[\bar{\mathrm{GLC}}][\bar{\mathrm{ATP}}]^2 + \delta_{[\bar{\mathrm{GLC}}]}J_{\mathrm{in}} \\
\frac{\partial[\bar{\mathrm{ATP}}]}{\partial\bar{t}} = & \frac{D_{[\mathrm{ATP}]}}{D_{[\mathrm{GLC}]}}\nabla^2[\bar{\mathrm{ATP}}] - 2\mathcal{K}_{\mathrm{HXK}}\beta_{[\bar{\mathrm{ATP}}]}[\bar{\mathrm{GLC}}][\bar{\mathrm{ATP}}]^2 + 2\mathcal{K}_{\mathrm{PYRK}}\gamma_{[\bar{\mathrm{ATP}}]}[\bar{\mathrm{ADP}}]^2[\bar{\mathrm{GLY}}] \\
& +28\mathcal{K}_{\mathrm{Mito}}\xi_{[\bar{\mathrm{ATP}}]}[\bar{\mathrm{PYR}}][\bar{\mathrm{ADP}}]^{28} - \mathcal{K}_{\mathrm{act}}\tau_{[\bar{\mathrm{ATP}}]}[\bar{\mathrm{ATP}}] \\
\frac{\partial[\bar{\mathrm{ADP}}]}{\partial\bar{t}} = & \frac{D_{[\mathrm{ADP}]}}{D_{[\mathrm{GLC}]}}\nabla^2[\bar{\mathrm{ADP}}] + 2\mathcal{K}_{\mathrm{HXK}}\beta_{[\bar{\mathrm{ADP}}]}[\bar{\mathrm{GLC}}][\bar{\mathrm{ATP}}]^2 - 2\mathcal{K}_{\mathrm{PYRK}}\gamma_{[\bar{\mathrm{ATP}}]}[\bar{\mathrm{ADP}}]^2[\bar{\mathrm{GLY}}] \\
& +\mathcal{K}_{Act}\tau_{[\bar{\mathrm{ADP}}]}[\bar{\mathrm{ATP}}] - 28\mathcal{K}_{\mathrm{Mito}}\xi_{[\bar{\mathrm{ADP}}]}[\bar{\mathrm{PYR}}][\bar{\mathrm{ADP}}]^{28} \\
\frac{\partial[\bar{\mathrm{GLY}}]}{\partial\bar{t}} = & \frac{D_{[\mathrm{GLY}]}}{D_{[\mathrm{GLC}]}}\nabla^2[\bar{\mathrm{GLY}}] + 2\mathcal{K}_{\mathrm{HXK}}\beta_{[\bar{\mathrm{GLY}}]}[\bar{\mathrm{GLC}}][\bar{\mathrm{ATP}}]^2 - \mathcal{K}_{\mathrm{PYRK}}\gamma_{[\bar{\mathrm{ATP}}]}[\bar{\mathrm{ADP}}]^2[\bar{\mathrm{GLY}}] \\
\frac{\partial[\bar{\mathrm{PYR}}]}{\partial\bar{t}} = & \frac{D_{[\mathrm{PYR}]}}{D_{[\mathrm{GLC}]}}\nabla^2[\bar{\mathrm{PYR}}] + \mathcal{K}_{\mathrm{PYRK}}\gamma_{[\bar{\mathrm{ATP}}]}[\bar{\mathrm{ADP}}]^2[\bar{\mathrm{GLY}}] - \mathcal{K}_{\mathrm{LDH}}\mu_{[\bar{\mathrm{PYR}}]}[\bar{\mathrm{PYR}}] \\
& -\mathcal{K}_{\mathrm{Mito}}\xi_{[\bar{\mathrm{PYR}}]}[\bar{\mathrm{PYR}}][\bar{\mathrm{ADP}}]^{28} \\
\frac{\partial[\bar{\mathrm{LAC}}]}{\partial\bar{t}} = & \frac{D_{[\mathrm{LAC}]}}{D_{[\mathrm{GLC}]}}\nabla^2[\bar{\mathrm{LAC}}] + \mathcal{K}_{\mathrm{LDH}}\mu_{[\bar{\mathrm{LAC}}]}[\bar{\mathrm{PYR}}] - \eta_{[\bar{\mathrm{LAC}}]}[\bar{\mathrm{LAC}}]
\end{cases}
\tag{1}
$$

where the dimensionless coefficients are shown in the Table 2:

| Dimensionless coefficients | | | | | |
|---|---|---|---|---|---|
| HXK | PYRK | LDH | Mito | act | |
| $\beta_{[\mathrm{G\bar{L}C}]} = t_c A_{\mathsf{tot}}^2$ | | | | | $\delta_{[\mathrm{G\bar{L}C}]} = \frac{t_c}{\alpha}$ |
| $\beta_{[\mathrm{A\bar{T}P}]} = t_c \alpha A_{\mathsf{tot}}$ | $\gamma_{[\mathrm{A\bar{T}P}]} = A_{\mathsf{tot}} \alpha t_c$ | | $\xi_{[\mathrm{A\bar{T}P}]} = t_c \alpha A_{\mathsf{tot}}^{27}$ | $\tau_{[\mathrm{A\bar{T}P}]} = t_c$ | |
| $\beta_{[\mathrm{A\bar{D}P}]} = t_c \alpha A_{\mathsf{tot}}$ | $\gamma_{[\mathrm{A\bar{D}P}]} = A_{\mathsf{tot}} \alpha t_c$ | | $\xi_{[\mathrm{A\bar{D}P}]} = t_c \alpha A_{\mathsf{tot}}^{27}$ | $\tau_{[\mathrm{A\bar{T}P}]} = t_c$ | |
| $\beta_{[\mathrm{G\bar{L}Y}]} = t_c A_{\mathsf{tot}}^2$ | $\gamma_{[\mathrm{G\bar{L}Y}]} = A_{\mathsf{tot}}^2 t_c$ | | | | |
| | $\gamma_{[\mathrm{P\bar{Y}R}]} = A_{\mathsf{tot}}^2 t_c$ | $\mu_{[\mathrm{P\bar{Y}R}]} = t_c$ | $\xi_{[\mathrm{P\bar{Y}R}]} = t_c A_{\mathsf{tot}}^{28}$ | | |
| | | $\mu_{[\mathrm{L\bar{A}C}]} = t_c$ | | | $\eta_{[\mathrm{L\bar{A}C}]} = \eta t_c$ |

Table 2: The table present the dimensionless coefficients for the dimensionless system 1.

In this way, we define a dimensionless system that depends only on the dimensionless volume of the cell.

### Supplementary Note 6: Detail on numerical methods for 2D simulations.

The 2D experiments were solved using standard finite element method (FEM) implemented in Python with the open source finite element solver DOLFIN from FEniCS [1,2]. The domains are explicitly meshed using the package *mshr*, which generate a finite element mesh that conforms to the boundary of the domains. The solution of the weak problem is defined on the space of piece wise Lagrange finite elements of degree one. We solve the non-linear equation using a Newton-Raphson scheme, where the Jacobian is calculated automatically by the automatic differentation capabilities of UFL [3]. The linear system at each time step of the Newton-Raphson algorithm is solved using standard linear solvers from the *PETSc* library. For further details [4].

### Supplementary Note 7: Details on numerical methods for 3D simulations.

To solve the system on the 3D astrocytic domains, we use a cut finite element [5] approach able to deal with the complexity of the cellular geometry, as shown in [4]. The difference between FEM and CutFEM lies in the fact that FEM requires to generate a mesh conformed to the boundary of the domains. This can be a difficult task and CutFEM removes this need. In CutFEM, we describe the boundaries implicitly through a level set function [6,7] that can be extracted from images. In particular, the level set function $\Phi$ is a scalar function that has negative values inside the domain, positive outside and zero on the boundary of the object. We obtained the level set, $\Phi$, of the final segmented image $f$ solving the following system in the domain $B$ which is a three dimensional box where we have mapped the image $f$:

$$\begin{cases} -\epsilon^2 \triangle \Phi + \Phi = f \quad \text{in}\, B \\ \nabla \Phi \cdot n = 0 \quad \text{on}\, \partial B \end{cases}$$

where $\epsilon$ is a smoothing parameter that we set to $0.001$. This step smooth the boundary of the cells to reduce the mesh size and related computational complexity respectively. In Supplementary Fig. 1, we show the regular background mesh of finite element covering the two astrocytic domains. To solve the weak formulation of the RDS using CutFEM, we define the fictitious domain as all the cells of the background mesh that have a non-zero intersection with the cellular domain (Supplementary Fig. 3). We apply a ghost-penalty stabilisation term [8] to all the edges that are intersected by the interface and all the edges that connect the intersected cells with the interior of the cellular domain. These stabilisation terms extend the solution from the physical domain $\Omega$ onto the fictitious domain. This is of fundamental importance to ensure the stability and accuracy of the numerical solution as the ghost-penalty stabilisation prevents ill-conditioning of the system matrices in case intersected cells contain very little of the cellular domain [5]. In this case, the linear system arising in the numerical experiments are solved using a direct (*MUMPS*) solver.

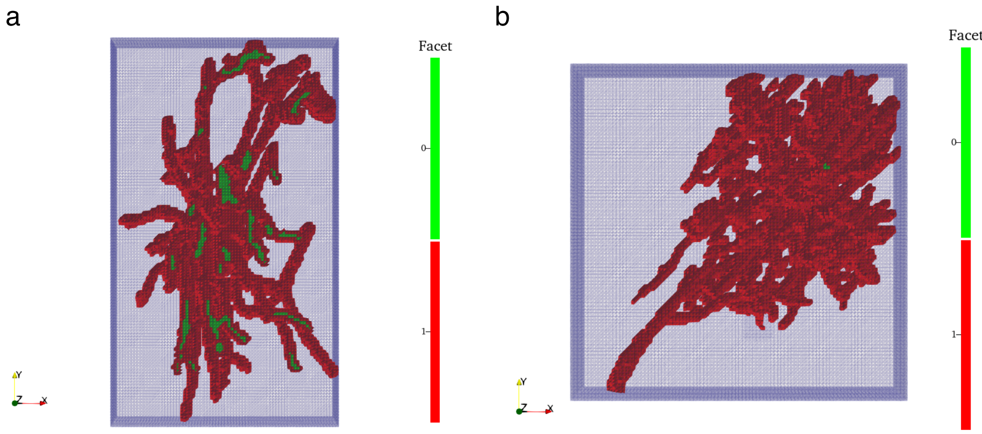


**Figure 3 Supplementary Fig.3: Fictitious domain and facet markers for the two astrocytic morphologies.** Fictitious domain for the two astrocytic shape **a** control and **b** reactive, defined as the non-zero intersection of the finite elements with the cellular domain. We apply the stabilization term to the facets marked in red.

For further reading we address the reader to our previous work [4] and alternative proposed for enriched FEM [9–12].

**Supplementary Note 8: Numerical Parameters.**

The parameters used for the 2D experiments are presented in Supplementary Table 3 and for the 3D experiments in Supplementary Table 4. The penalty parameter $\gamma$ used to ensure stability on the cut cells is set to $0.1$. Convergence study were done extensively in the 2 dimensional experiments.

| 2D Numerical parameters | | | | |
|---|---|---|---|---|
| geometry | # cells | # dofs | $\max$ cell diameter | $\Delta t$ |
| circle | 88768 | 44856 * | $0.93$ | $0.25$ |
| star | 51046 | 26164 * | $0.77$ | $0.25$ |
| rectangle | 25298 | 13207* | 0.33 | 0.17 |

Table 3: The table presents the numerical parameters used in the 2D experiments for the three domains: circle, star and rectangle. We show the number of cells in the finite element mesh, the number of degrees of freedom (dofs), the maximum diameter of the finite element cell and the time step used for the time discretisation. * In the table the number of dofs refers to one subspace. The total number of dofs for all the six subspaces is 269136 for the circle, 156984 for the star and 79242 for the rectangle.

| 3D Numerical parameters | | | | | |
|---|---|---|---|---|---|
| astrocyte | # cells in bg mesh | # cells on $\Omega$ | # dofs | $\max$ cell diameter | $\Delta t$ |
| control | 1000008 | 158938 | 37742 | 0.022 | 0.08 |
| reactive | 1301760 | 162579 | 39143 | 0.018 | 0.026 |

Table 4: The table presents the numerical parameters used in the 3D experiments for the control and reactive astrocytes. We show the number of cells in the background finite element mesh, the number of cell covering the astrocyte domains $\Omega$, the number of degrees of freedom (dofs), the maximum diameter of the finite element cell and the time step used for the time discretisation.
